# Performance of blood-based biomarkers for human circadian pacemaker phase: Training sets matter as much as feature selection methods

**DOI:** 10.1101/2025.05.21.655317

**Authors:** Carla S. Möller-Levet, Simon N Archer, Derk-Jan Dijk

## Abstract

Biomarkers are valuable tools in a wide range of human health areas including circadian medicine. Valid, low burden, multivariate molecular approaches to assess circadian phase at scale in people living and working in the real-world hold promise for translating basic circadian knowledge to practical applications. However, standards for the development and evaluation of these circadian biomarkers have not yet been established, even though several publications report such biomarkers and claim that the methods are universal. Here we present a basic exploration of some of the determinants and confounds of blood-based biomarker development for SCN phase by reanalysing publicly available data sets. We compare performance of biomarkers based on three feature selection methods: Partial Least Squares Regression, ZeitZeiger and Elastic Net, as well as performance of a standard set of clock genes. We explore the effects of training sample size and the impact of the experimental protocols from which training samples are drawn and on which performance is tested. Approaches based on small sample sizes used for training are prone to poor performance due to overfitting. Performance to some extent depends on the feature selection method, but at least as much on the experimental conditions from which the biomarker training samples were drawn. Performance of biomarkers developed under baseline conditions do not necessarily translate to protocols that mimic real world scenarios such as shiftwork in which sleep may be restricted or desynchronised from endogenous circadian SCN phase. The molecular features selected by the various approaches to develop biomarkers for SCN phase show very little overlap, although the processes associated with these features have common themes with response to steroid hormones, i.e. cortisol being the most prominent. Overall, the findings indicate that establishment of circadian biomarkers should be guided by established biomarker development concepts and foundational principles of human circadian biology.

## Introduction

Circadian rhythmicity and disturbances thereof play a significant role in health and disease (Cederroth et al., 2019). Prime examples are circadian rhythm sleep disorders (Meyer et al., 2022), cancer and cancer therapeutics (Karaboue et al., 2024; Zeng et al., 2024), mental health disorders (Meyer et al., 2024), neurological disorders (Khan et al., 2018) and cardio-metabolic disorders (Thosar et al., 2018; Arab et al., 2024; Marhefkova et al., 2024). To further quantify the contribution of circadian rhythmicity to health and disease and elucidate the underlying mechanisms, methods to accurately assess circadian rhythmicity that can be implemented at scale need to be developed (Mullington et al., 2016; Dijk and Duffy, 2020; Pundir et al., 2022). In recent years several new methods to quantify circadian rhythmicity have been developed (for a summary see Dijk and Duffy, 2020). Among the methods under development, biomarkers based on panels of transcripts, metabolites and proteomes that only require one or a few samples to quantify circadian rhythmicity, rather than a time series as is the case for classical markers such as melatonin, cortisol and core body temperature, have received much attention. As for biomarkers in general (Biomarkers Definitions Working, 2001), these circadian biomarkers do not measure circadian rhythmicity directly but are used as an indicator or surrogate marker of circadian rhythmicity. It is often claimed that these biomarkers may be used to identify abnormal circadian rhythmicity and the response to interventions but for now these novel biomarkers are not yet implemented in research or clinical practise.

Given that in mammals circadian rhythmicity is present in all tissues and cells (Mure et al., 2018; Patton and Hastings, 2023), a first specification in the development of biomarkers relates to the organ, tissue, brain area, cell type or organelle that is targeted by the biomarker. A second area of consideration relates to the aspect of rhythmicity that is to be monitored by the biomarker. Quantifying circadian rhythmicity can focus on traditional parameters such as period, phase or amplitude (Kuhlman et al., 2018) or can consider alternative descriptors such as circadian clock function (Vlachou et al., 2024) which may be defined as an appropriate constellation of core elements of circadian rhythmicity, e.g., products of clock genes.

In recent literature the phrase circadian rhythmicity is often used in a rather loose manner to mean 24-h rhythmicity as observed, i.e., diurnal rhythmicity. However, in the quest for the identification of the endogenous self-sustaining oscillators driving 24-h rhythmicity the distinction between endogenous and evoked contributions to overt ‘circadian’ rhythmicity has played a key role (Aschoff, 1960; Kuhlman et al., 2018). Rhythmicity in the peripheral transcriptome is driven in part directly by circadian processes but also by the sleep-wake cycle and associated rhythmic factors such as feeding and light exposure (Archer et al., 2014; Jan et al., 2024). In many applications ‘circadian’ rhythmicity is quantified, and biomarkers are to be implemented under conditions in which environmental and behavioural factors will contribute to rhythmicity (e.g., Bowman et al., 2021). Although it is unclear whether it is overt rhythmicity or the endogenous circadian component that is most relevant to, for example, health outcomes, it is relevant to specify whether the biomarker targets endogenous components or merely attempts to capture aspects of rhythmicity as it is observed.

Biomarker identification can not only be applied as a simple marker but may also provide clues about mechanisms underlying the association between the process targeted by the biomarker and outcomes, e.g., how does circadian disruption lead to negative health outcomes (Biomarkers Definitions Working, 2001; Pourali et al., 2024). In this context it is relevant that the features that comprise the biomarker and the associated biological processes are characterised.

Circadian biomarkers consisting of panels of features selected from samples of whole blood or specific cell types, hair follicles and specific layers of skin have featured in the recent literature (Kasukawa et al., 2012; Laing et al., 2017; Braun et al., 2018; Wittenbrink et al., 2018; Wu et al., 2018; Braun et al., 2019; Lee et al., 2020; Wu et al., 2020; Cogswell et al., 2021; Woelders et al., 2023). Although in several cases the purpose of the biomarker has been stated explicitly, in other cases it was not clear whether the purpose was to assess rhythmicity of the tissue from which the sample was taken or rhythmicity in the human circadian pacemaker, i.e., the SCN. It is nevertheless almost always implied that the biomarker may be useful to assess circadian phase in circadian rhythm disorders, disorders related to shift work, ageing and other conditions in which circadian rhythmicity is considered relevant. However, the training sets used to construct the biomarker rarely mimicked these conditions and standard markers of the SCN such as plasma melatonin were not always used to evaluate the performance of these biomarkers. Consequently, it is not clear to what extent the reported performance of these biomarkers reflects SCN phase and translates to conditions such as circadian rhythm sleep disorders or shift work.

The biomarkers in these recent studies are based on the selection of a set of features but the selection methods vary from a-priori selection of a set of clock genes, selection from a set of rhythmic genes (e.g., Zeitzeiger) or unbiased methods (e.g., Partial Least Squares Regression or Elastic Net) which select features from all available features without any prespecified characteristic. To what extent the feature selection method influences the composition of the feature panel used to quantify circadian rhythmicity and the performance of these biomarkers across conditions remains an open question.

Here, we will focus on blood-based biomarkers for the assessment of SCN phase as indexed by what is considered the gold standard marker of circadian phase, i.e., melatonin phase derived from time series of blood or saliva samples. We focus on biomarkers for SCN phase because this pacemaker is assumed to synchronise oscillators throughout the body and plays a critical role in circadian rhythm sleep-wake disorders, shift work and jet lag disorders. The specific questions we address are:

1. To what extent do performance of biomarkers for SCN circadian phase depend on the size of the training set?
2. Is biomarker performance across conditions dependent on the algorithm and training sets used to identify the predictor sets?
3. Can biomarkers for circadian phase be used in conditions other than the conditions used to develop the biomarker?
4. Do predictor sets, along with their associated genes and processes, vary across the algorithms and conditions used for their development?

We address these questions by applying several feature selection methods to data collected in protocols in which the timing of the sleep-wake cycle, sleep history, posture and the light-dark cycle were varied and gold standard markers of SCN phase, i.e., melatonin, were available.

## Methods

### Data

All analyses were based on publicly available datasets from the NCBI Gene Expression Omnibus (accessions: GSE39445, GSE48113, GSE82113, GSE82114, and GSE253864). Detailed information about these studies, including participant demographics, procedures and ethics approval, can be found in the associated publications (Moller-Levet et al., 2013; Archer et al., 2014; Laing et al., 2017; Archer et al., 2024). For each sample, circadian phase was assessed relative to Dim Light Melatonin Onset (DLMO), as described in Laing et al., 2017 and Bonmati-Carrion et al., 2024.

### Microarray data processing

For each of the five GEO datasets, raw data in the form of Feature Extraction text files were downloaded from the GEO repository.

For each training set, log2 mRNA abundance values were quantile-normalized using the limma R package (v 3.58.1), and the empirical quantiles values were recorded. Non-control technically replicated probes (mRNA abundance features) were averaged, and control probes were removed from the dataset, resulting in a total of 41,619 transcripts. Transcript values were then z-scored across samples, with mean and standard deviation values recorded for each transcript.

For each validation set, the log2 mRNA values were quantile-normalized using the same reference array of empirical quantiles from the training set. This approach maintains the same empirical distribution between the training and validation data and thereby helps minimise distributional differences that may arise from technical variability. Non-control technically replicated probes (mRNA abundance features) were averaged, and control probes were removed. Transcript values were z-scored using the mean and standard deviation obtained from the training set. To integrate all datasets, we used the probes present in the microarray platform (GEO accession no. GPL15331). This ensured consistency across datasets.

### Training and validation set construction

Samples from the Forced Desynchrony (Archer et al., 2014) and Sleep Restriction and Total Sleep Deprivation studies (Moller-Levet et al., 2013) were used to generate five different training sets: sleeping in phase (IP), sleeping out of phase (OP), sufficient sleep (SS), insufficient sleep (IS), and a set that includes an equal number of samples from the four conditions (All). The model resulting from each training set was used to predict the circadian phase of samples from eleven different validation sets: IP, OP, SS, IS, all, and the six conditions from the ESA Head Down Tilt (HDT) Bed Rest study, which consisted of 2 weeks of baseline data collection (BDC), 60 days of constant HDT, and 2 weeks of recovery (R). Time series samples were collected twice during baseline (BDC1, BDC2), three times during HDT (HDT1, HDT2, HDT3), and once during recovery (R) (Archer et al., 2024). A diagram illustrating this procedure is presented in Supplemental Figure 1.

Each training set was generated by randomly selecting a predefined number of samples, allowing overlap in participant between the training and validation sets. To account for the variation introduced by random selection, 20 training sets were created for each condition (IP, OP, SS, IS, and All), resulting in 20 distinct models per condition. Although training sets with unique (non-repeating) participants would yield statistically independent samples, such an approach reduces the representation of inter-individual variability – a critical consideration given the highly individual nature of blood transcriptome profiles (Battle et al., 2014).

Nevertheless, we also tested a non-overlapping sampling strategy and observed no substantial improvement in model performance, supporting the use of overlapping sets to better capture biological diversity.

### Modelling methods description and implementation

In the main analyses, we compared three machine learning algorithms: Partial Least Squares Regression (PLSR) (Boulesteix & Strimmer, 2007), ZeitZeiger (ZZ) (Hughey et al., 2016), and Elastic Net (EN) (Zou & Hastie, 2005). PLSR and EN are widely used machine learning algorithms and have been applied in the development of biomarkers for circadian phase (Laing et al., 2017; Braun et al., 2018), whereas ZZ was specifically designed for this purpose. We also completed a less extensive evaluation of the performance of biomarkers for circadian phase based on a defined Set of Clock Genes.

#### Partial Least Squares Regression (PLSR)

PLSR creates uncorrelated latent factors from the original variables and then performs linear regression on these factors. These latent factors are linear combinations of the original predictors, summarizing the information from all predictors. The weights of each predictor in forming these latent factors are estimated by maximizing the covariance between the predictor variables and the response variable. Through this process, PLSR implicitly selects features, with the most relevant predictors for the model being associated with higher weights. The use of latent factors allows PLSR to addresses high dimensionality and multicollinearity.

The *plsr* function from the pls R package (v 2.8-3) was used to perform PLSR with the SIMPLS algorithm. The number of predictor probes and the number of latent factors were set to 100 and 5, respectively, based on Laing et al., 2017. The top 100 predictor probes are selected based on the highest sum of absolute weights across the 5 latent factors. During model fitting, the regression coefficients for these 100 predictors are estimated. The regression is performed in Cartesian coordinates, where each training circadian angle is decomposed into its cosine and sine components, resulting in two dependent variables (the sine and cosine components) and two regression coefficients for each predictor. The predicted circadian phase is calculated from the predicted Cartesian coordinates (see Laing et al., 2017 for details).

#### ZeitZeiger (ZZ)

The Zeitzeiger algorithm (Hughey et al., 2016) models time-dependent feature patterns using splines to capture how features vary over time. This time-structured data is then converted into a discretized data matrix and subjected to penalized matrix decomposition, which isolates the variation in features associated with time. The resulting sparse principal components (SPCs) are linear combinations of a select few features, driven by ***l1*** regularization. This regularization ensures that the principal components are sparse, meaning only the most important features contribute to them. This reduces the risk of overfitting. After decomposition, the training data is projected from the original feature space into the new SPC space using the loadings (weights) of each feature in the SPCs. Splines are then reapplied to estimate the time-dependent patterns of these SPCs. For time prediction of a test sample, Zeitzeiger uses maximum-likelihood estimation, relying on the SPC values of the test sample and the previously estimated time-dependent patterns of the top SPCs from the training data. As in PLSR and EN, the features with the highest loadings on the SPCs are considered the most important for the model.

The *zeitzeigerFit* and *zeitzeigerSpc* functions from the Zeitzeiger R package (v 2.1.3) were used to fit periodic smoothing splines to the measurements of each feature as a function of time and to the calculate the sparce principal components of time-dependant variation. The *zeitzeigerPredict* function was used for time prediction in test samples. The main parameters of Zeitzeiger, sumabsv (***l1*** regularization) and nSPC (number of SPCs), were set to 4 and 3, respectively, based on Laing et al., 2017.

#### Elastic Net (EN) Regression

EN combines the strengths of Lasso and Ridge regression by performing linear regression with both ***l1*** and ***l2*** penalties on the model’s coefficients. The ***l1*** penalty promotes sparsity by shrinking some coefficients to zero, effectively selecting the most important predictors, while the ***l2*** penalty stabilizes the model by grouping correlated predictors together. This dual regularization enables Elastic Net to perform feature selection and regularization simultaneously, addressing high dimensionality and multicollinearity while ensuring that the most relevant predictors are retained in the model.

The function *glmnet* from the glmnet R package (v 4.1-8) was used to perform Elastic Net regression. The alpha parameter, which determines the balance between Lasso and Ridge regularization, was set to 0.5, reflecting a combination of both methods (where alpha = 1 is pure Lasso, and alpha = 0 is pure Ridge). The lambda parameter, which controls the strength of the penalty, was set to the value that minimizes the mean cross-validated error obtained using the *cv.glmnet* function (Laing et al., 2017) and the training and validation sets (lambda = 0.03280596). Like PLSR, the regression is performed in Cartesian coordinates, where each training circadian angle is decomposed into its cosine and sine components, resulting in two dependent variables (the sine and cosine components) and two regression coefficients for each predictor. The predicted circadian phase is calculated from the predicted Cartesian coordinates (see Laing et al., 2017 for details).

The three modelling methods involve hyperparameters that need to be optimised during model development. For practicality and ease of comparison, we fixed these parameters based on the hyperparameter optimization reported in Laing et al. (2017) for both PLSR and ZZ and determined the optimal lambda for EN using the same training and validation datasets. However, for completeness, we also examined how the optimal hyperparameters varied across training conditions for PLSR and ZZ and assessed the impact on performance of using optimized versus fixed lambda values for EN.

### Evaluation of prediction performance

We assessed model performance using three metrics:

1. Mean Absolute Error (MAE): This metric measures the average magnitude of absolute prediction errors, i.e. ignoring the direction of the error, providing an indication of accuracy which is often used.
2. Mean Error: This metric with its standard deviation measures the average of the prediction errors and their distribution. The average indicates the bias.
3. R^2^ (Coefficient of Determination): This metric compares the variance of prediction errors to the variance in observed circadian phases. R^2^ is calculated as described in Laing et al., 2017. R^2^ reflects the proportion of variance explained by the model, with vales closer to 1 indicating better performance. Negative R2 values indicate poor performance, where the model is worse than a horizontal line fit.

Note: Even though MAE is an often-used metric it is not without problems because it cannot distinguish between cases in which all errors are positive, all errors are negative and cases in which errors are both positive and negative. In other words, it cannot assess the average bias. The Mean error and its standard deviation in principle contains all this information but requires a slightly better understanding of the metrics of distributions and is therefore often not used.

All angles (*θ)* were converted to Cartesian coordinates (*cosine θ, sine θ*) to perform operations such as calculating the directional mean and angle differences. For further details, refer to Laing et al., 2017. The standard deviation of angles was calculated using the sd.circular function from the circular R package (v 0.5-0).

The effect of different variables (e.g., number of samples in the training set, modelling method, training set, validation set) and their interactions on circadian phase prediction performance was assessed via ANOVA using the aov R function.

The Compact Letter Display (CLD), used to visualise the significance of contrasts, was generated using the multcompLetters4 function from the multcompView R package (v0.1-10), based on the results of Tukey’s pairwise comparisons (TukeyHSD R function). This analysis was conducted following a two-way ANOVA for each modelling method, which assessed the main effects of both the training and validation sets, as well as their interaction on performance.

### Predictor transcripts

For a specific training set, overlapping predictors were defined as those that appeared in at least 50% of the models generated (i.e., in at least 10 out of 20 models). For models generated using PLSR and EN, weights for each predictor transcript were calculated as the square root of the sum of squares of the regression coefficients for the sine and cosine terms. In models generated with ZZ, weights for each predictor transcript were calculated as the square root of the sum of squares of the SPC loadings. In all cases, weights were averaged across runs. To summarise at the gene level, weights from all transcripts belonging to the same gene were averaged.

The number of overlapping predictors was obtained under five different training conditions and three different training set sizes, producing 15 values per modelling method. The coefficient of variation (CV, i.e., standard deviation/mean) of the 15 values was then computed to assess consistency. A lower CV indicates that the overlap of predictors in random re-runs of the same dataset is similar across different training conditions and sample sizes, meaning the method’s consistency in feature selection is stable across training sets with different characteristics. Conversely, a higher CV reflects varying degrees of consistency in feature selection across training sets with different characteristics.

The intersection of overlapping predictors across training sets and modelling methods was evaluated using ‘upset plots’, a data visualization technique that displays intersecting sets and their relationships. Upset plots were generated using upset function from the ComplexUpset (v 1.4.0) R package and customised using the ggplot2 (v 3.5.0) and patchwork (1.2.0) R packages.

In addition to standard feature selection methods, we also tested PLSR using core clock genes as a fixed predefined list of predictors. Specifically, we utilized a set of 24 classic clock genes identified in (Moller-Levet et al., 2022), which were mapped to 504 microarray probes and used as predictors in the generation of the PLSR models.

### Effect of training set sample selection strategy on biomarker performance

Twenty training sets of 100 randomly selected samples were generated for each experimental condition. The models resulting from each training set were used to predict the circadian phase of samples in the validation sets. We evaluated the effect of the sample selection strategy on biomarker performance by comparing the mean MAE values (mean across the 20 models per training condition) of models generated using: (A) randomly selected training samples, and (B) training and validation sets with no overlapping participants. We also examined the phase distribution of the samples in the selected training sets and evaluated MAE performance as a function of circadian phase by grouping validation samples into 30° phase bins across the circadian cycle.

### Impact of number of Samples used for Training

A training set was created by randomly selecting ***n_s_*** samples from the total ***N*** samples of condition ***X****_1-5_* (IP, OP, SS, IS, all). The ***n_s_*** values used were 60, 100 and 140. The validation set included the remaining (***N***-***n_s_***) samples from condition ***X***, along with all samples from the other conditions. A predictive model was developed using each training set and then used to estimate circadian phase in the validation set. The predicted and observed circadian phase were compared, and the performance was calculated independently for each experimental condition in the validation set. To account for the variation introduced by random selection, 20 training sets were created per condition. A three-way ANOVA was then conducted to estimate the effects of the method, the training set condition, the number of samples in the training set (***n_s_***), and their interactions.

### Effect of modelling (i.e. feature selection) method on biomarker performance

A training set was created by randomly selecting 140 samples from the total ***N*** samples of condition ***X****_1-5_*. The validation set included the remaining (***N***-140) samples from condition ***X***, along with all samples from the other conditions. A predictive model was developed using each training set and then used to estimate circadian phase in the validation set. The predicted and observed circadian phase were compared, and the performance was calculated independently for each experimental condition in the validation set. To account for the variation introduced by random selection, 20 training sets were created per condition. A two-way ANOVA was conducted to estimate the effects of the method, the training set condition, and their interactions on MAE. Additionally, a one-way ANOVA was performed within each training set to evaluate the effect of method, followed by Tukey’s pairwise comparisons to identify differences in MAE.

### Impact of Training Set on Performance under Conditions different from the training set

A training set was created by randomly selecting 140 samples from the total *N* samples of condition *X*. The validation set included the remaining (*N*-140) samples from condition *X*, along with all samples from other conditions. A predictive model was developed using each training set and then used to estimate circadian phase in the validation set. The predicted and observed circadian phase were compared, and the performance was calculated independently for each experimental condition in the validation set. To account for the variation introduced by random selection, 20 training sets were created per condition. A three-way ANOVA was then conducted to estimate the effects of modelling method, training set, validation set and their interactions.

### Impact of Training Set and Feature Selection Method on selected Features and Associated Processes

Gene ontology (GO) enrichment analyses were performed using Webgestalt 2024 (webgestalt.org). The GO databases used were ‘Biological Process noRedundant’ and ‘Molecular Function noRedundant’. The Agilent human wholegenome 4×44k v2 was used as the reference set, which was the microarray platform used for each study in the analyses presented here.

## RESULTS

### Available data

Across the 10 data collection sessions (Fig 1) 1,419 transcriptomic samples were used for the analyses. These samples were collected in a total of 82 participants. Details on the demographics can be found in the original publications. Circadian melatonin phase, derived from blood or saliva, was assigned to each of these samples as described previously.

**Figure 1.**
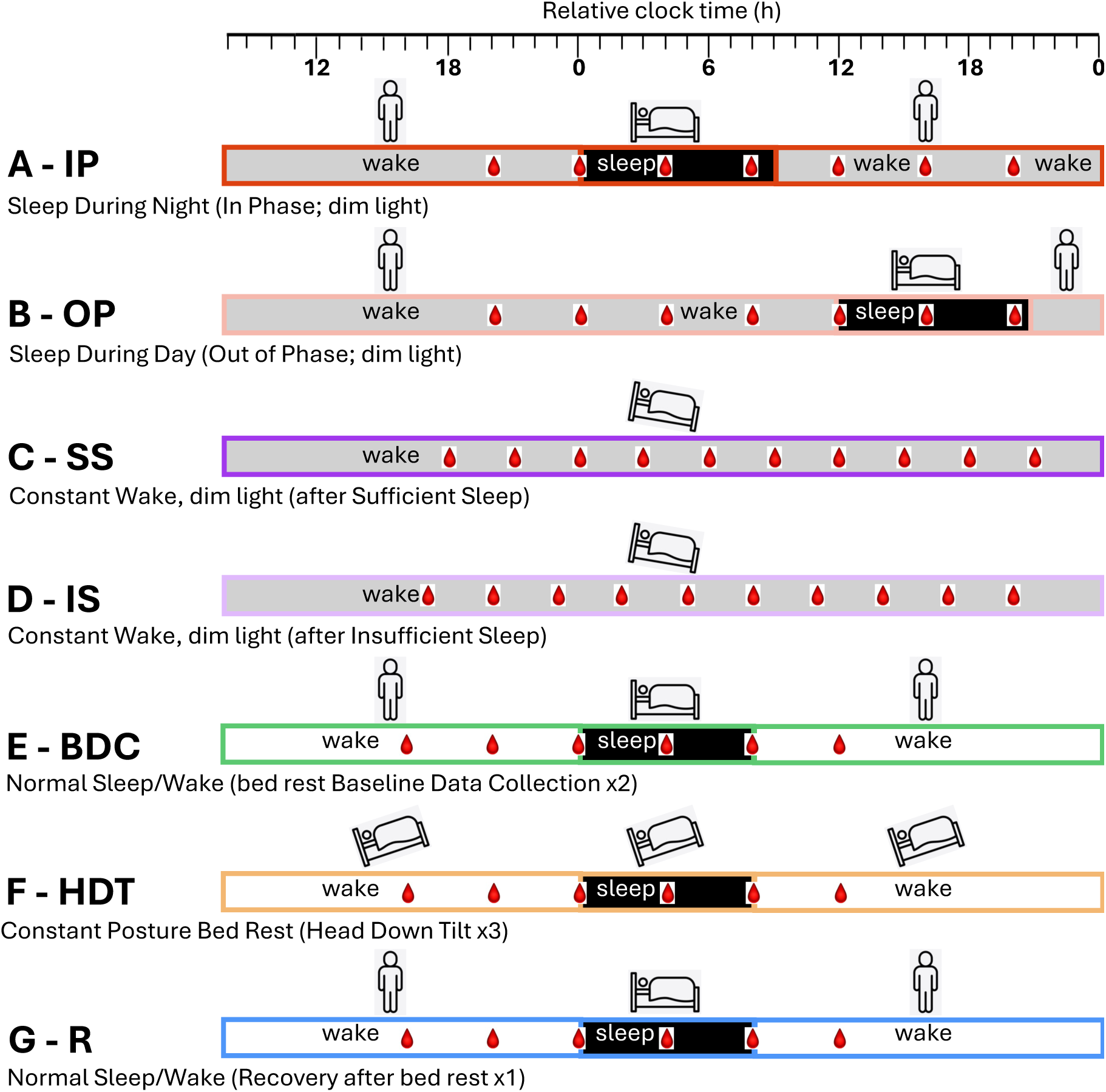
Schematic of the ten sleep/wake conditions utilised in phase biomarker evaluation. Horizontal bars represent the different sleep/wake conditions: **A)** sleeping in phase with the circadian clock during the night (black) and wake in dim light (gray); **B**) sleeping out of phase during the day with wake in dim light; **C)** constant wake in constant routine (CR) conditions (dim light, semi recumbent, regular frequent meals, etc.) after one week of sufficient sleep (average 8.5h); **D)** constant wake during CR after one week of insufficient sleep (average 5.7h); **E)** normal sleep/wake (sleep in darkness, wake in normal room light) during two 24h sessions in the baseline data collection period (2 weeks) of a bed rest study; **F)** sleep/wake during three 24h sessions within a 60-day period of constant posture bed rest at a -6 degree head down tilt; **G)** normal sleep/wake during one 24h session of a 2-week recovery period after constant bed rest. For each sleep/wake condition, bedtime is set to 0h (relative clock time). During each sleep/wake condition, blood sampling for transcriptomics is indicated by a red blood drop. In all conditions, circadian phase was assessed from melatonin rhythms in blood (A-D) or saliva (E-G). A and B Archer et al., 2014, C and D Moller-Levet et al., 2013, E – F Archer et al., 2024.

### Effect of training set sample selection strategy on biomarker performance

We evaluated the effect of the construction of the training and validation sets on biomarker performance by comparing the MAE values of models generated using: (A) randomly selected training samples, and (B) training and test set with no overlapping participants (Figure 2). The Pearson correlation (r) of MAE values between strategies A and B is r=0.75, p=2.2×10⁻¹⁶, with n=1,200 (5 training conditions × 20 training sets × 4 validation conditions × 3 modelling methods). The range of differences in mean MAE values between strategies A and B (A-B across training conditions and modelling methods) spans from 0 to 0.2 hours (12 minutes) (see Figure 2).

**Figure 2.**
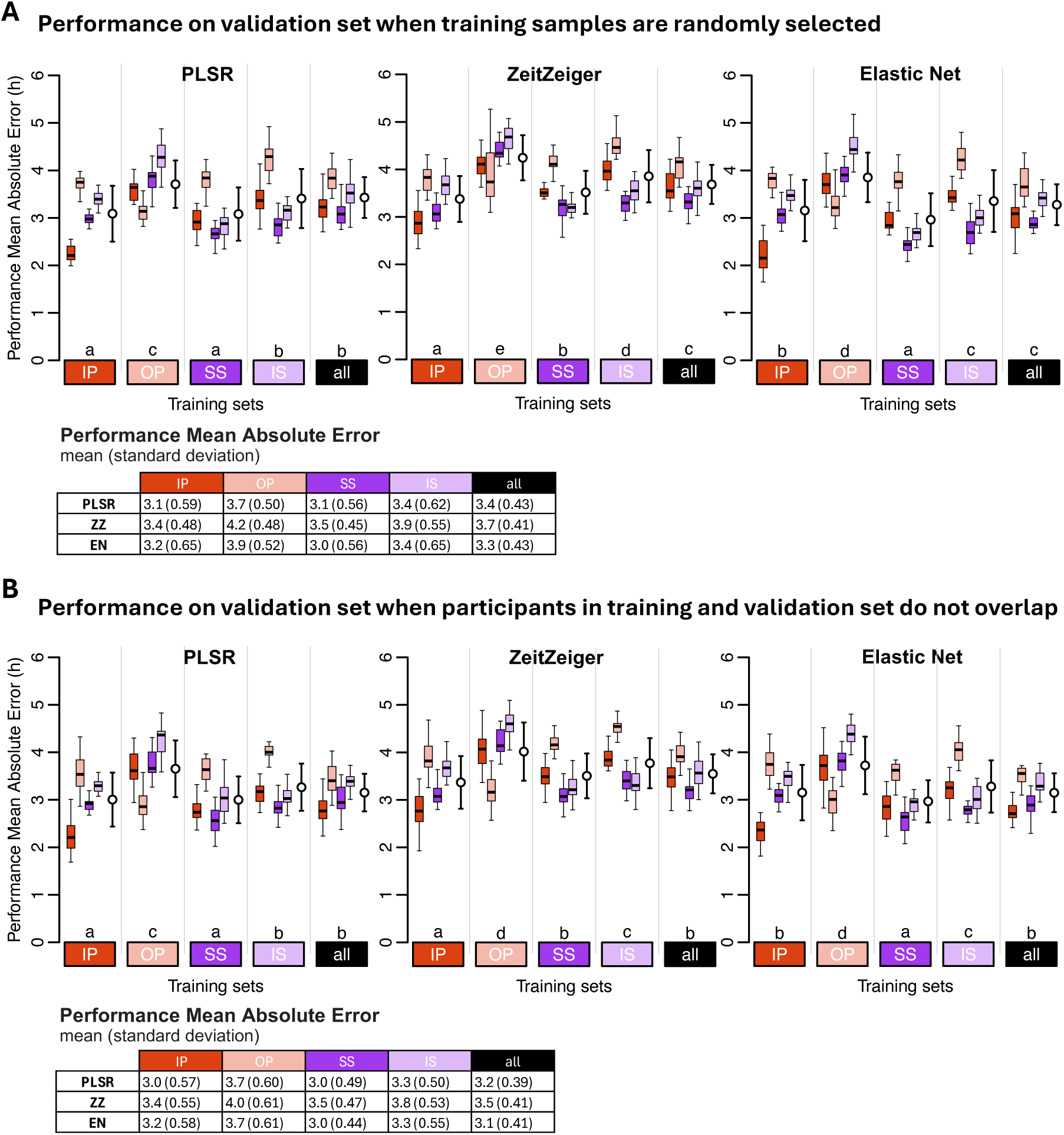
Effect of sample selection strategy on biomarker performance. Performance on (**A**) validation set, when training samples are randomly selected, and (**B**) on validation set when participants in training and validation do not overlap. Performance results obtained for PLSR (left), ZeitZeiger (center), and Elastic Net (right). Twenty training sets of 100 samples were generated for each experimental condition. Each coloured boxplot represents 20 mean absolute error (MAE) values for each condition of the validation set, with outliers omitted for clarity. White open circles with error bars indicate the mean and standard deviation of all 80 validation values per training set (20 sets x 4 validation conditions). Letters above the training sets represent the Compact Letter Display (CLD) which summarizes the results of pairwise comparisons between different training sets. Groups sharing the same letter are not significantly different, while groups with different letters are significantly different (p < 0.05). The p values are determined by Tukey’s test conducted on a two-way ANOVA of training and validation sets within each method. Tables present the mean and standard deviation of 80 MAE values per training condition (20 training sets x 4 validation conditions). In all panels, training sets are depicted as coloured rectangles on the horizontal axis. The training condition “all” includes an equal number of samples from each of the four experimental conditions.

We also examined the phase distribution of the samples in the selected training sets and evaluated MAE performance as a function of circadian phase by grouping validation samples into 30° phase bins across the circadian cycle (supplemental figure 2). Both sample selection strategies produce training sets with samples that adequately span the circadian cycle. Some over-representation of specific time points is observed in both strategies, reflecting the time series structure, which results in one or two samples being collected at the same clock time, 24 hours apart. The resulting MAE patterns are consistent across both selection strategies, with low variation across bins. In supplemental figure 2, panel C demonstrates strong significant correlations (all BH p<0.05) between the phase-binned MAE values from panels A (randomly selected training samples) and B (randomly selected training samples from non-overlapping participants), confirming that prediction performance patterns are highly similar regardless of whether participant overlap in the training and validation sets is permitted.

Together, these analyses support the robustness of our findings across different training sample selection strategies and confirm that the observed model performance patterns are not driven by participant overlap.

### Effect of size of training sets on biomarker performance

We first investigated the effect of size of the training set on model performance on the training set. the performance on the training set decreased with increasing sample size and this for all methods and all training sets (reduction in performance sample size 140-60: 0.65,0.8, and 0.1 hours for PLSR, ZZ and EN, respectively) (Fig 3A and 3C). The Mean Absolute Error (MAE) performance of the biomarkers developed on samples from protocols, A-IP, B-OP, C-SS or D-IS (Fig 1) was calculated with performance being assessed by comparing the predicted phase to the observed phase of all remaining samples in these four protocols. The performance on the validation set (Fig 3B and 3D) was poorer than the performance on the training sets (and this for all three methods and sample sizes. The average difference (averaged over the three sample sizes) between training and validation performance was 2.4, 1.8 and 3.3 hours for PLSR, ZZ and EN respectively. The performance on the validation set significantly increased when the training set increased from 60, to 100 and 140 (Fig 3B and 3D). This pattern was observed for all training sets and for all three methods (Partial Least Squares Regression, Zeitzeiger and Elastic Net). The absolute error on the validation set when the training set sample size was increased from 60 to 140, decreased by 1.1, 0.6, and 1.1 hours for PLSR, ZZ and EN, respectively. The performance of the models on the validation set also depended on the protocol used to develop the biomarker (i.e., the training set). The three-way ANOVA for MAE used to assess the effects of the method, training set, number of samples in the training set, and their interactions yielded significant main effects for the three variables (all p<2×10^-16^) and their interactions (all p<0.001). The effect of training set size on improvement of performance on the validation set was not directly related to the performance on the training set. The effect size of training sets on performance, based on Mean Error and R^2^, follows similar trends and can be found in Tables S1 and S2, respectively.

**Figure 3.**
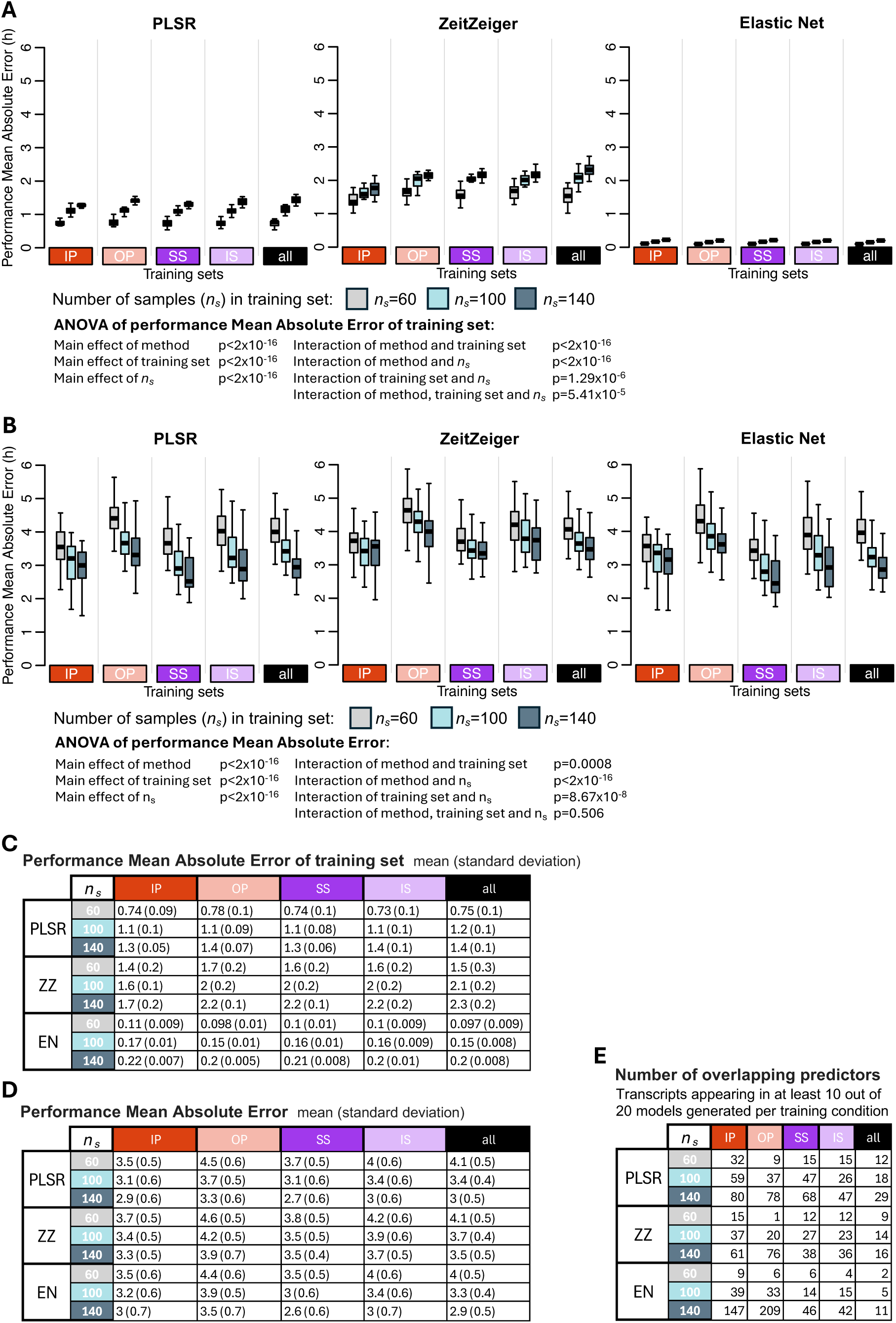
Effect of number of samples in the training set on biomarker performance. **A** Performance results of training set, where the same samples are used for training and validation, obtained for the three modelling algorithms: PLSR (left), ZeitZeiger (centre), and Elastic Net (right). Each box plot represents the distribution of 20 mean absolute error (MAE) values for each condition of the validation set, totalling 80 values (20 sets x 4 conditions), with outliers omitted for clarity. **B** Performance of validation set, where different samples are used for training and validation. Each box plot represents the distribution of 20 MAE values, with outliers omitted for clarity. **C** Mean and standard deviation of 80 MAE values per training condition (20 training sets x 4 validation conditions) shown in panel **A**. **D** Mean and standard deviation of 20 MAE values per training condition (20 training sets, where the same samples are used for training and validation) shown in panel **B**. **E** Number of overlapping predictors, defined as transcripts appearing in at least 10 out of the 20 models generated per training set. In each panel, results were obtained for *n_s_* values of 60 (gray box plots), 100 (light blue box plots) and 140 (dark blue box plots), and training sets are represented by coloured rectangles on the horizontal axis. The training condition “all” includes an equal number of samples from each of the four experimental conditions. IP = sleeping in phase during the night, OP = sleeping out of phase during the day, SS = constant wake after one week of sufficient sleep, IS = constant wake after one week of insufficient sleep.

Figure 3E shows that the number of overlapping predictors (defined as those appearing in at least 10 out of 20 models per training condition) selected by the three methods (PLSR, ZZ, EN) varied under different sample sizes (60, 100, 140) and across training conditions (IP, OP, SS, IS, and all). The largest coefficient of variation (CV) of the number of overlapping predictors across all sample sizes and conditions was observed for EN (1.50), while the smallest CV was observed for PLSR (0.63). Within each method, the largest CV consistently occurred for the OP condition, while the smallest CV was consistently observed for the “all” condition across all methods.

Since the number of predictors can be predetermined for PLSR, we also evaluated the interaction between the number of predictor transcripts and the size of the training set on PLSR performance. This was done by comparing the MAE values of models generated using different numbers of predictor transcripts (*n_t_* = 50, 100 and 150) and varying numbers of training samples (*n_s_* = 60, 100, 140) (Fig 4). Visual inspection of Fig 4 suggests that the effect of number of predictors is small. A three-way ANOVA for MAE was conducted to assess the statistical significance of effects of the training set, number of samples in the training set (***n_s_***), number of predictor transcripts (***n_t_***), and their interactions. The analysis revealed significant main effects for all three factors (p<2×10^-16^, p<2×10^-16^ and p=1.84×10^-5^, respectively), as well as a significant interaction between training set and ***n_s_*** (p=7.17×10^-8^). However, the interactions between training set and ***n_t_***, ***n_s_*** and ***n_t_***, and the three-way interaction among training set, ***n_s_*** and ***n_t_*** were not significant (p>0.05 for all). These results confirm that while performance varies with predictor count, the methods remain robust even with smaller training sets.

**Figure 4.**
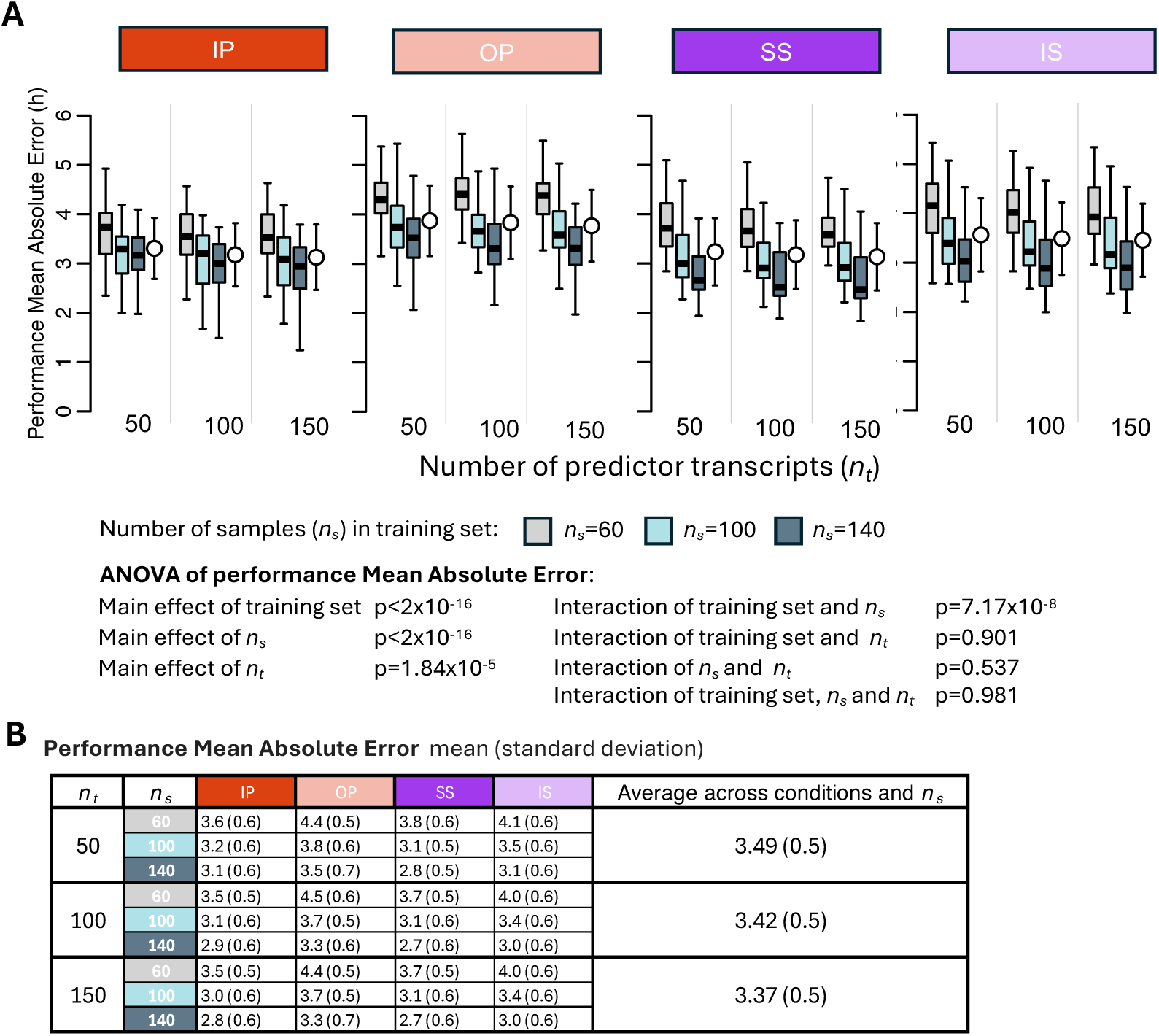
Effect of number of predictor transcripts on PLSR performance. **A** Performance results obtained for PLSR using different numbers of predictor transcripts (*n_t_* = 50, 100 and 150) and different numbers of samples in the training set (*n_s_* = 60, 100, 140; gray, light blue, and dark blue box plots, respectively). Each box plot represents the distribution of 20 mean absolute error (MAE) values for each condition of the validation set, totalling 80 values (20 sets x 4 conditions), with outliers omitted for clarity. **B** Mean and standard deviation of 80 MAE values per training condition (20 training sets x 4 validation conditions) shown in panel **A**. Training sets are represented by coloured rectangles on the horizontal axis. IP = sleeping in phase during the night, OP = sleeping out of phase during the day, SS = constant wake after one week of sufficient sleep, IS = constant wake after one week of insufficient sleep.

#### Effect of Biomarker Construction method on performance across a range of protocols

We evaluated the performance of different biomarker construction methods (PLSR, ZZ, and EN) across a range of training conditions (IP, OP, SS, IS, and all, where the “all” condition includes an equal number of samples from all conditions).

The R^2^ values - which quantify how well the predicted phases align with the observed phases, with higher values indicating better agreement - were similar for PLSR and EN across training conditions and significantly higher than those observed for ZZ (Fig 5A). The performance of each method varied depending on the training condition, with OP showing the lowest R^2^ values and the ‘all’ condition achieving the highest R^2^ values.

**Figure 5.**
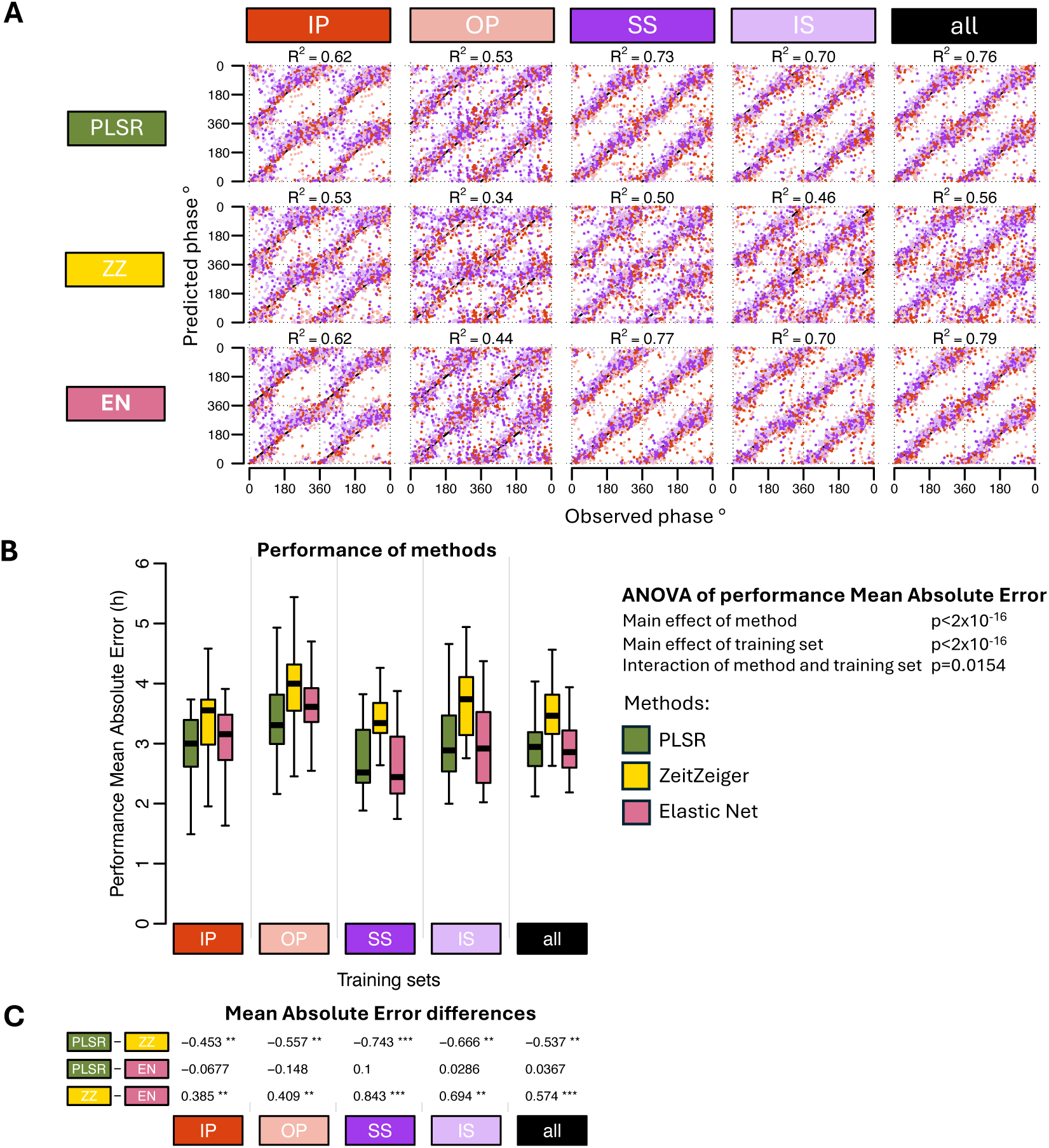
Effect of modelling method on biomarker performance. **A** Predicted circadian phase of a blood sample vs. observed melatonin phase. The predicted circadian phase (averaged across *n* runs, where *n* is the number of runs out of 20 in which a sample was included in the validation set) is plotted against the observed melatonin phase for each validation sample, color-coded by condition. All training sets consisted of 140 samples. Circadian phase 0 corresponds to the onset of the melatonin rhythm, which occurred on average at 23:13 ± 10 min across all conditions. The data are double-plotted, and the line of unity represents perfect prediction. Results are shown for three modelling algorithms - PLSR, ZeitZeiger (ZZ), and Elastic Net (EN) (rows) - across five different training sets (columns). The R2 quantifies how well the predicted angles align with the observed angles, with higher values indicating better agreement. **B**. Performance results obtained using PLSR (green box plots), ZeitZeiger (yellow box plots), and Elastic-net (pink box plots). Twenty training sets of 140 samples were generated for each experimental condition. Each box plot represents the distribution of 20 mean absolute error (MAE) values for each condition of the validation set, totalling 80 values (20 sets x 4 conditions), with outliers omitted for clarity. A two-way ANOVA for MAE revealed significant main effects of both method and training set (both p < 2×10⁻¹⁶), and a significant interaction between them (p = 0.0154). **C** Results of a one-way ANOVA conducted within each training set to assess the effect of the method on MAE. Tukey’s pairwise differences in MAE and the associated p-values (*** p < 1×10⁻¹², ** p < 0.01, and * p < 0.05) are shown, with rows indicating pairwise comparisons and columns representing the experimental conditions. Training sets are represented by coloured rectangles on the horizontal axes. The training condition “all” includes an equal number of samples from each of the four experimental conditions. IP = sleeping in phase during the night, OP = sleeping out of phase during the day, SS = constant wake after one week of sufficient sleep, IS = constant wake after one week of insufficient sleep.

Overall, the performance measured as mean absolute error was rather large ranging from approximately 3-4 hours when assessed on samples drawn from all conditions (Fig 5B). Comparing the performance of biomarkers developed from samples on a variety of protocols and validated across all conditions demonstrated a significant effect of Method (p<2×10^-16^). Zeitzeiger performed significantly poorer than both PLSR and EN (Fig 5C) and this for all protocols used for biomarker development. In this context it is relevant to mention that ZZ’s performance was also worst on the training sets (Fig 3A and 3C). The effect of biomarker construction method on performance, based on Mean Error and R^2^, follows similar trends and can be found in Tables S1 and S2, respectively.

#### Effect of training set and validation set and their interaction on performance

The protocol used as training set and the protocol used to test the performance both had a significant effect on performance, as measured by Mean Absolute Error. Additionally, the interaction between the training and validation set was also significant (ANOVA all p < 2×10⁻¹⁶) (Fig 6). For all three methods and all training protocols, the performance of biomarkers was best when assessed on the remaining samples of the training set. For example, PLSR, Zeitzeiger and Elastic Net biomarkers developed on samples from participants sleeping at night (IP) performed better on the remaining samples of IP than on samples from any of the other conditions.

**Figure 6.**
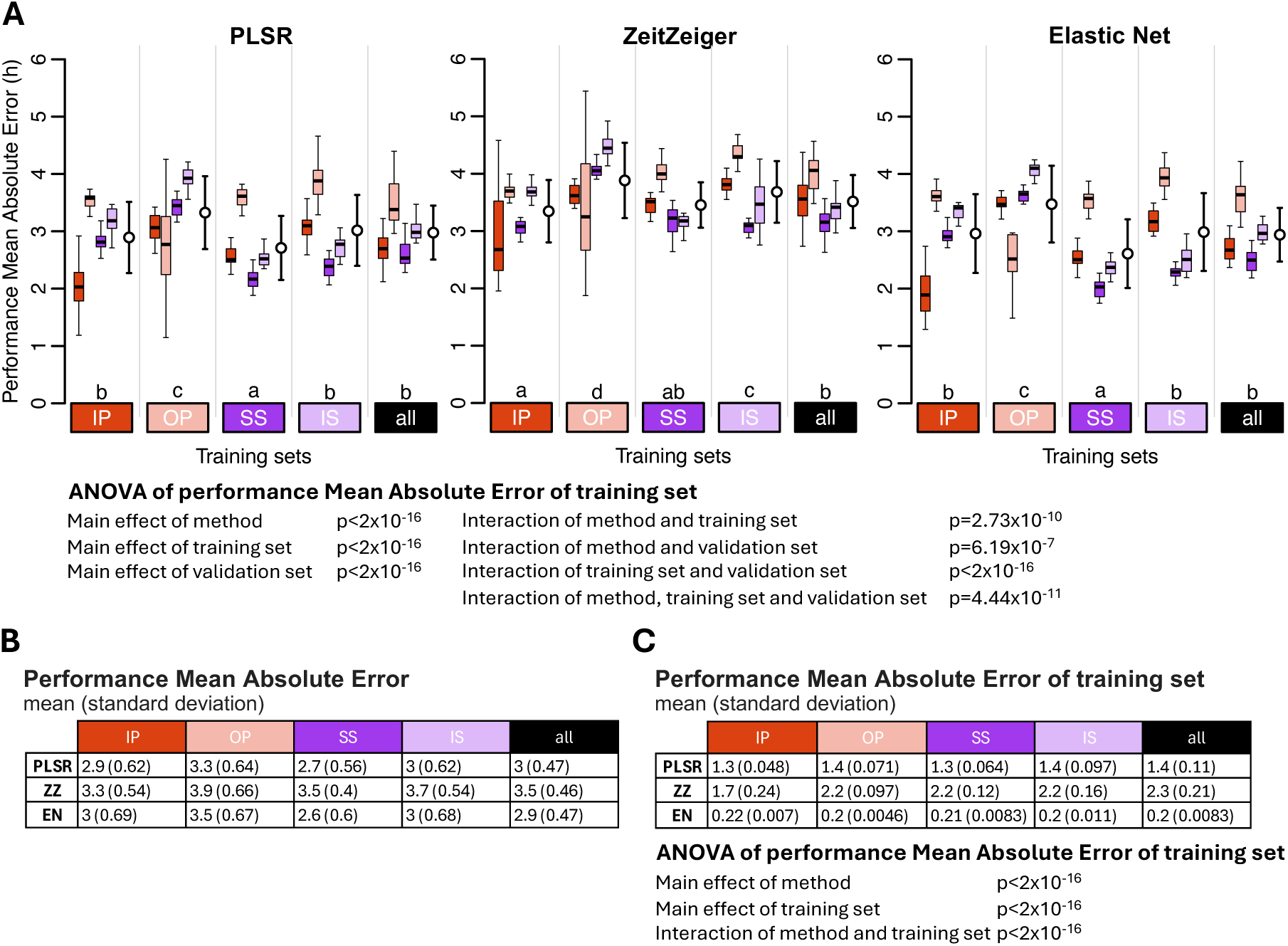
Effect of training set and validation set on biomarker performance for the three feature selection methods. **A** Performance results obtained for PLSR (left), ZeitZeiger (center), and Elastic Net (right). Twenty training sets of 140 samples were generated for each experimental condition. Each coloured box plot represents the distribution of 20 mean absolute error (MAE) values for each condition of the validation set, with outliers omitted for clarity. White open circles with error bars indicate the mean and standard deviation of all 80 validation values per training set (20 sets x 4 conditions). The three-way ANOVA for MAE revealed significant main effects of method, training set, and validation set (all p < 2×10⁻¹⁶) and significant two-way interactions: interaction between method and training set (p =2.73×10⁻¹^0^), interaction between method and validation set (p=6.19×10^-7^), and interaction between training set and validation set (p < 2×10⁻¹⁶). The three-way interaction of method, training set, and validation set was also significant (p=4.44×10^-11^). Letters above the training sets represent the Compact Letter Display (CLD) which summarizes the results of pairwise comparisons between different training sets. Groups sharing the same letter are not significantly different, while groups with different letters are significantly different (p < 0.05). The p values are determined by Tukey’s test conducted on a two-way ANOVA of training and validation sets within each method. **B** Mean and standard deviation of 80 MAE values per training condition (20 training sets x 4 validation conditions) shown in panel **A**. **C** Mean and standard deviation of 20 MAE values per training condition (20 training sets per condition), where the same samples are used for both training and validation. The two-way ANOVA for MAE of the training set showed significant effects of method and training set, and a significant interaction between them (all p < 2×10⁻¹⁶). In all panels, training sets are depicted as coloured rectangles on the horizontal axis. The training condition “all” includes an equal number of samples from each of the four experimental conditions. IP = sleeping in phase during the night, OP = sleeping out of phase during the day, SS = constant wake after one week of sufficient sleep, IS = constant wake after one week of insufficient sleep.

Likewise, biomarkers developed on samples from participants who were awake for approximately 40 h after a week of sufficient sleep (SS) performed better on the remaining samples collected in this condition than on samples collected in any of the other conditions and this so for all three selection methods. Of particular interest is the performance of biomarkers developed on samples collected from participants on a normal sleep-wake cycle, i.e., being awake during the day and sleeping at night (IP), when applied to samples collected in participants sleeping during the day (OP). For all three methods the performance was then rather low. Biomarkers developed on the sleeping out of phase condition performed rather variably across the conditions and even when assessed on the remaining out of phase samples and this for all three methods. Biomarkers developed on samples collected during constant routine conditions (SS and IS) and in particular the biomarkers developed based on samples collect during CR conditions when participants were well rested (SS) upon entry into the constant routing performed consistently better across all conditions. Similar statements apply to biomarkers developed on samples derived from a combination of all protocols, and the same effects were observed when performance was calculated using the Mean Error and R^2^ (see Tables S1 and S2).

#### Performance on a completely different dataset-protocol

When applied to a data set collected under conditions of baseline, bedrest with head down tilt and recovery from this challenge, biomarkers developed from samples collected while donors were sleeping in phase or when in constant routine conditions performed best (Fig 7 and Table S3). Biomarkers developed from samples while participants slept out of phase performed worst. Across methods, Zeitzeiger performed worst and PLSR performed best. For PLSR and EN the performance on this data set was similar to their performance on combined IP, OP, SS, IS protocols. Within the bedrest data set, performance was lowest on samples collected during the recovery phase. The biomarker performance based on Mean Error and R^2^ follows similar trends and can be found in Tables S1 and S2, respectively.

**Figure 7.**
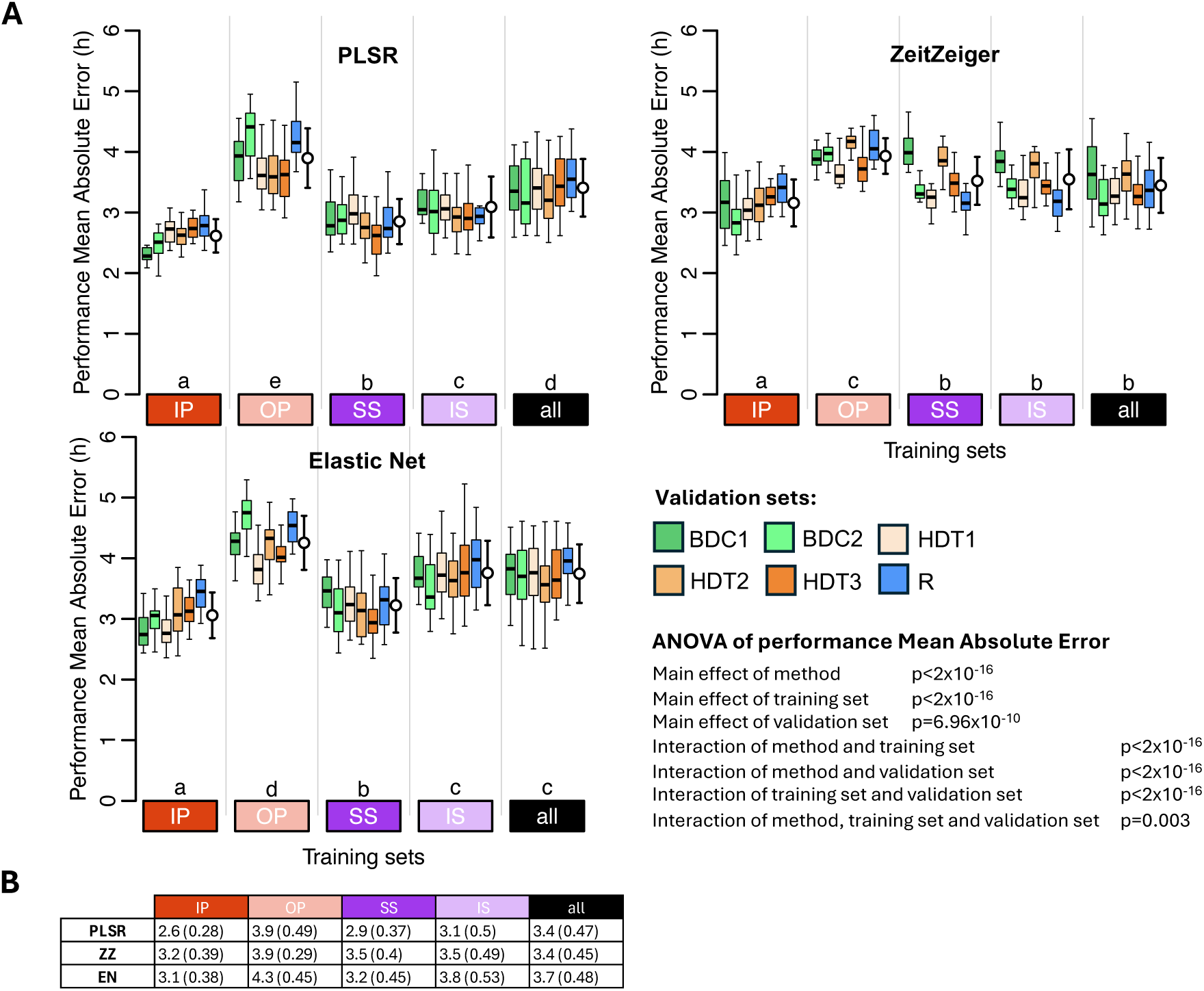
Performance of biomarkers on an independent data set collected in a bed rest protocol. A. Performance results obtained for PLSR (left), ZeitZeiger (center), and Elastic Net (right). Twenty training sets of 140 samples were generated for each experimental condition. Each box plot represents the distribution of 20 mean absolute error (MAE) values for each condition in the validation set, with outliers omitted for clarity. White open circles with error bars indicate the mean and standard deviation of all 120 validation values per training set (20 sets x 6 conditions). A three-way ANOVA for MEA revealed significant main effects of method (p < 2×10⁻¹⁶), training set (p < 2×10⁻¹⁶), and validation set (p=6.96×10-10). Significant interactions were found between method and training set, training set and validation set, and method and validation set (all p < 2×10⁻¹⁶). The three-way interaction among method, training set, and validation set was also significant (p = 0.003). The Compact letter display (CLD) summarises the results of pairwise comparisons between different training sets, as determined by Tukey’s test conducted on the two-way ANOVA of training and validation sets within each method. B Mean and standard deviation of 120 MAE values per training condition (20 training sets x 6 validation conditions) shown in panel A. In all panels, training sets are depicted as coloured rectangles on the horizontal axis. The training condition “all” includes an equal number of samples from each of the four experimental conditions. BDC1 = bed rest baseline data collection 1, BDC2 = bed rest baseline data collection 2, HDT1 = constant bed rest head down tilt 1, HDT2 = constant bed rest head down tilt 2, HDT3 = constant bed rest head down tilt 3, R = recovery after constant bed rest.

#### Performance based on PLSR applied to a pre-defined set of clock genes

PLSR was applied to a fixed set of 24 clock genes in five training sets, and performance was assessed on five evaluation sets (Supplemental Fig 3). In this approach, the features remained constant across training sets, i.e. they are fixed to the 24 clock genes, but their coefficients (weight) varied. Overall, the performance of this approach was poorer than when PLSR was used to select features (and determine their coefficients) (compare Supplemental Fig 3B & 3C to Fig 6A & 6B). For IP and SS and similar to the other methods, performance was best when assessed on the remaining samples of the training set. Performance on samples collected during desynchrony of the sleep-wake cycle and melatonin rhythm was in general poor. Performance based on Mean Error and R^2^ follows similar trends and can be found in Tables S1 and S2, respectively.

#### Model Hyperparameters selection

For both PLSR and ZZ, we used leave-one-participant-out cross-validation to identify the optimal combination of model parameters. For PLSR, this involved selecting the number of latent factors and predictors; for ZZ, it involved selecting the number of sparse principal components (nSPC) and the sumabsv regularization value. The PLSR results (supplemental figure 4) indicated that five latent factors and 100 predictors were optimal across all four training conditions (IP, OP, SS, IS), consistent with the fixed parameters used in the current study, and reported in Laing et al. (2017). In contrast, the optimal parameters for ZZ (supplemental figure 5) varied by condition (IP: nSPC = 2, sumabsv = 2.5; OP: nSPC = 3, sumabsv = 2.5; SS: nSPC = 4, sumabsv = 2; IS: nSPC = 2, sumabsv = 4). These results suggest that while PLSR hyperparameters are more robust across different datasets, the optimal ZZ parameters are more data-dependent. In the current study we fixed nSPC=3 and sumabsv=4 as reported in Laing et al. (2017).

While a grid search can be used for PLSR and ZeitZeiger due to their few and independent hyperparameters, for Elastic Net, we fixed alpha and optimized lambda using cross-validation along a regularization path. This method is more efficient because lambda is continuous, and its effect depends on the value of alpha. In this case we re-generated the models using optimised lambda values, rather than the fixed lambda used originally. The results of this optimisation (supplemental figure 6) show a very high consistency with the previous analysis: the correlation between the MAE values obtained with the fixed and optimised lambda across training sample sizes and training conditions is 0.99 (p=1.2×10^-12^, *n*= 15). The range of differences in mean MAE values between the two approaches (fixed lambda – optimized lambda) spans from –0.1 to 0.2 hours (from -6 to +12 minutes), indicating minimal impact on model performance relative to the differences derived from other variables.

#### Predictor sets vary with training set and method with little overlap

Figures 8 and 9 show upset plots with the intersection of overlapping predictors (defined as those appearing in at least 10 out of 20 models per training condition) across training conditions within each modelling method (Fig 8), and across methods within each sleep/wake condition (Fig 9). The intersection of overlapping predictors across training sets varies by method, with minimal overlap observed overall. For the four training conditions (IP, OP, SS, IS), the percentages of predictors unique to each condition are as follows: PLSR - 48.8%, 66.7%, 38.2%, and 42.6%; ZeitZeiger - 60.7%, 78.9%, 21.1%, and 19.4%; and Elastic Net - 87.8%, 92.3%, 56.5%, and 57.1%. For all three methods, the training set for ‘all’ conditions had the smallest number of overlapping predictors. This means that when the method had to train upon samples from all conditions, there was more variability in the predictors selected in each of the 20 runs, such that few predictors appeared in 50% of the runs. This observation is consistent with the finding that in general predictors were more specific for each sleep/wake condition.

**Figure 8.**
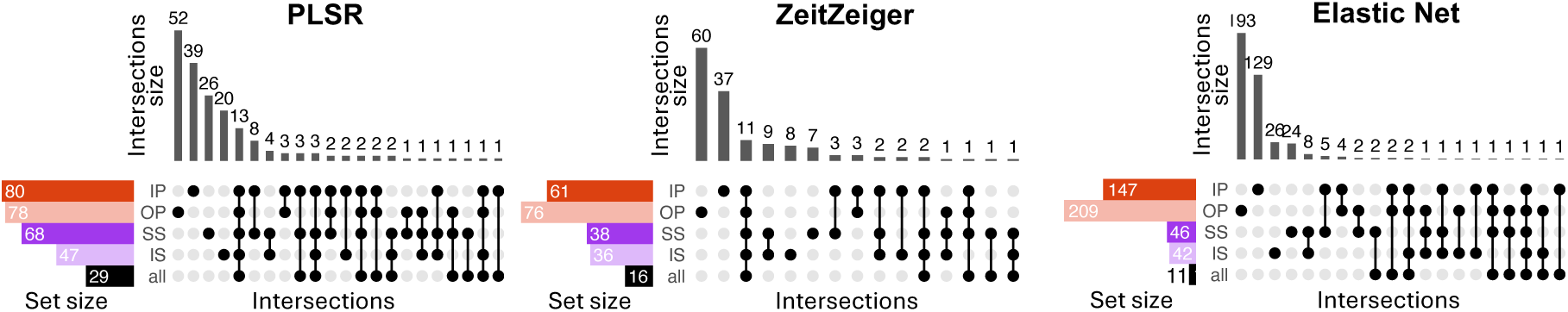
Intersection of overlapping predictors across training sets. Results for models generated using PLSR (left), Zeitzeiger (centre) and Elastic Net (right). Twenty training sets of 140 samples were generated for each experimental condition. Overlapping predictors were defined as those appearing in at least 10 out of the 20 models generated. In each panel, each row represents a set of overlapping predictors, while each column indicates a possible intersection of overlapping predictors across training conditions. Filled circles show which set is part of an intersection and the bar charts on the top show the number of elements in the intersection. IP = sleeping in phase during the night, OP = sleeping out of phase during the day, SS = constant wake after one week of sufficient sleep, IS = constant wake after one week of insufficient sleep. The condition “all” includes an equal number of samples from each of the four experimental conditions.

**Figure 9.**
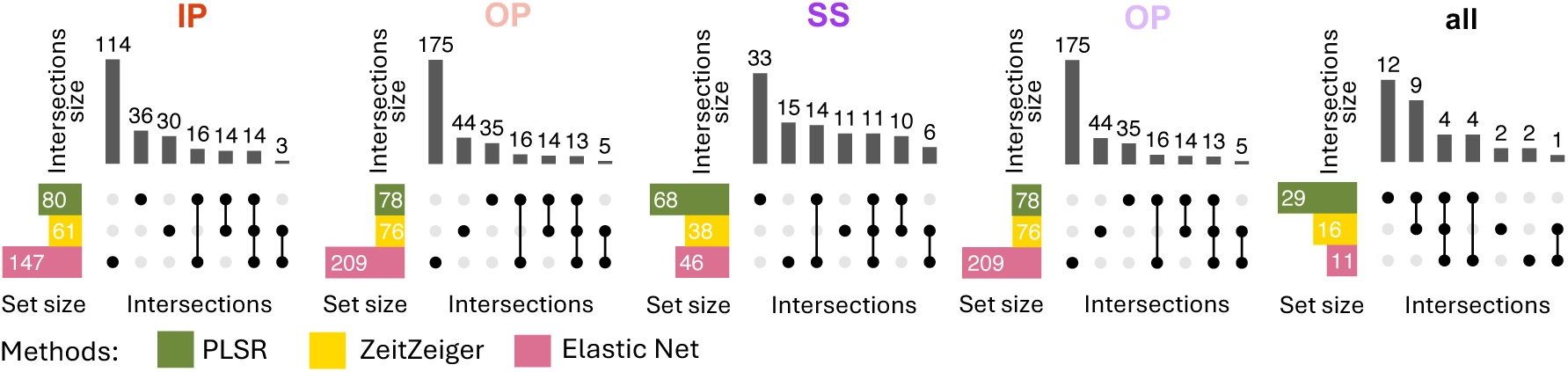
Intersection of overlapping predictors between methods. Results are displayed from left to right for models generated with samples from IP, OP, SS, IS, or all conditions combined. Twenty training sets of 140 samples were generated for each experimental condition. Overlapping predictors were defined as those appearing in at least 10 out of the 20 models generated. In each panel, each row represents a set of overlapping predictors, while each column indicates a possible intersection of overlapping predictors across methods (PLSR, Zeitzeiger and Elastic Net). Filled circles show which set is part of an intersection and the bar charts on the top show the number of elements in the intersection. IP = sleeping in phase during the night, OP = sleeping out of phase during the day, SS = constant wake after one week of sufficient sleep, IS = constant wake after one week of insufficient sleep. The condition “all” includes an equal number of samples from each of the four experimental conditions.

When comparing the intersection of overlapping predictors across methods for each training set condition (Fig 9) the same trend was observed, i.e., little overlap across methods.

Table 1 lists the overlap of genes coded by the training set predictor probes between the 3 training methods for each training set condition. *PER1* and *DDIT4* are predictors identified by all three methods for each training set condition, while *PER3* and *FKBP5* appeared in 4 conditions (IP, OP, SS, IS, and IP, OP, IS, ALL, respectively), and *TSPAN4* and *GPER* appeared in 2 (SS, IS, and IP, OP, respectively). All other genes were predictors across all 3 methods but only for single training sets. *PER1*, *PER3* and *NR1D2* are all core circadian clock genes, and *RBM3* is a temperature sensitive RNA binding protein that regulates the expression of clock genes. In this analysis, probes for *PER1* are predictors of circadian phase that are robust against sleep loss and mistimed sleep, unlike *Per2* which has been shown previously to be sleep/wake-driven (Hoekstra et al., 2021). In addition, *PER1*, *DDIT4* and *FKBP5* are induced within glucocorticoid signalling pathways and featured as robust phase predictors in our previous study (Laing et al., 2017).

**Table 1.**
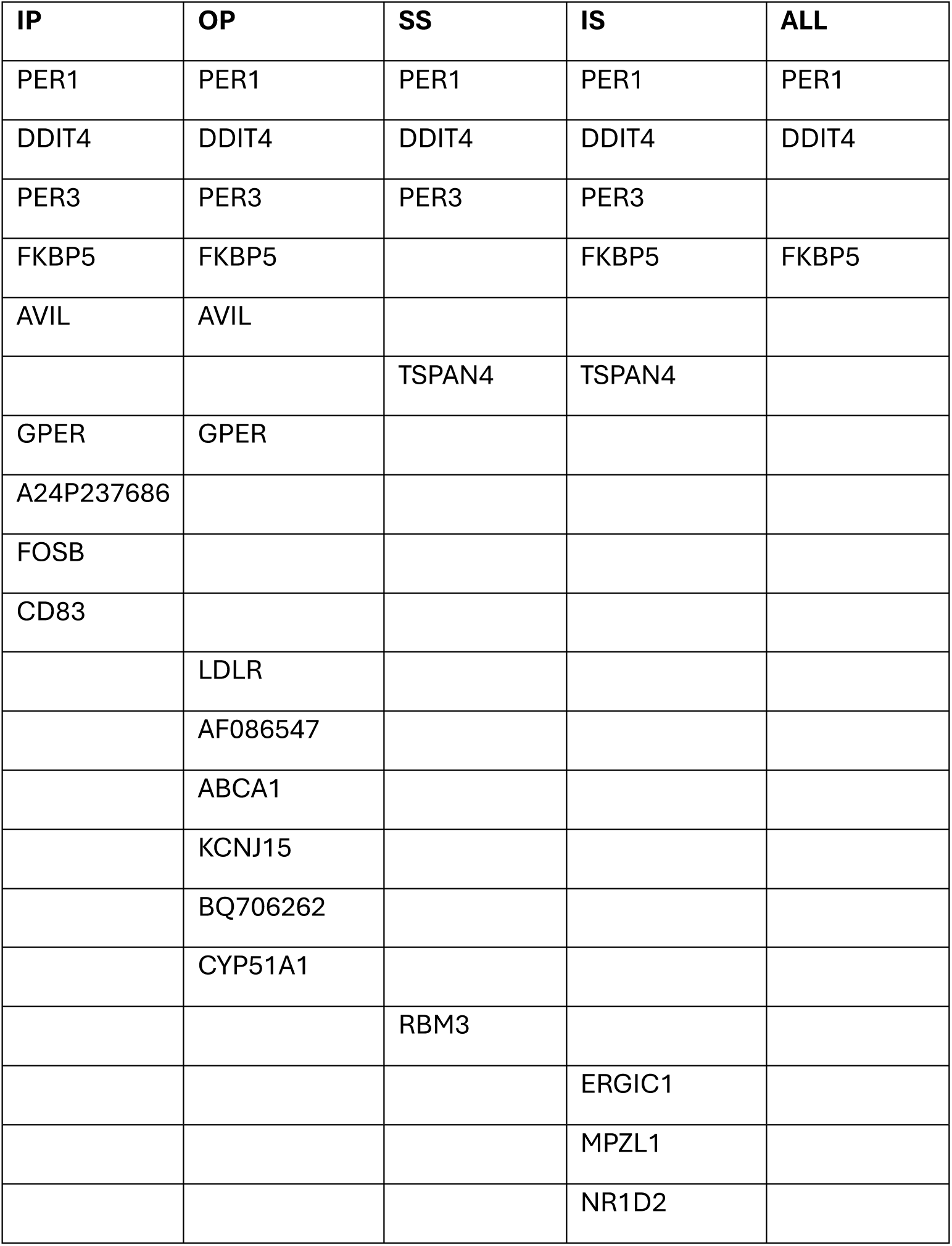
Overlap of genes coded by training set predictor probes between the 3 training methods for each training set condition.

| IP | OP | SS | IS | ALL |
| --- | --- | --- | --- | --- |
| PER1 | PER1 | PER1 | PER1 | PER1 |
| DDIT4 | DDIT4 | DDIT4 | DDIT4 | DDIT4 |
| PER3 | PER3 | PER3 | PER3 |  |
| FKBP5 | FKBP5 |  | FKBP5 | FKBP5 |
| AVIL | AVIL |  |  |  |
|  |  | TSPAN4 | TSPAN4 |  |
| GPER | GPER |  |  |  |
| A24P237686 |  |  |  |  |
| FOSB |  |  |  |  |
| CD83 |  |  |  |  |
|  | LDLR |  |  |  |
|  | AF086547 |  |  |  |
|  | ABCA1 |  |  |  |
|  | KCNJ15 |  |  |  |
|  | BQ706262 |  |  |  |
|  | CYP51A1 |  |  |  |
|  |  | RBM3 |  |  |
|  |  |  | ERGIC1 |  |
|  |  |  | MPZL1 |  |
|  |  |  | NR1D2 |  |

#### Biological processes and molecular functions are somewhat consistent across methods and training sets

Table 2 shows the top 5 gene ontology (GO) terms for biological processes and molecular functions enriched (P < 0.05) within the predictor probe sets for each training method and for each training set condition. GO terms that occurred across multiple methods and training sets are indicated in bold and colour-coded were identical. The results clearly show consistent enrichment for probes associated with ‘rhythmic process’, ‘rhythmic behaviour’, and ‘transcription coregulator binding’ across conditions for PLSR and EN (and to a lesser extent for ZZ). ‘Response to steroid hormone’ also featured within PLSR and ZZ training sets. These results confirm the contribution of clock genes and glucocorticoid signalling genes as predictor probes.

**Table 2.** Top 5 gene ontology terms for biological processes and molecular functions enriched within probe sets for each training method and for each training set condition.

|  | PLSR | ZZ | EN |
| --- | --- | --- | --- |
| IP | **response to steroid hormone**<br>sleep<br>response to ketone<br>fat cell differentiation<br>transcription coregulator binding | e-box binding<br>**response to steroid hormone**<br>response to radiation<br>response to organophosphorus<br>response to purine-containing compound | response to fatty acid<br>regulation of lipid metabolic process<br>steroid metabolic process<br>rhythmic process<br>regulation of small metabolic process |
| OP | rhythmic process<br>cellular response to lipoprotein stimulus<br>**response to steroid hormone**<br>response to nutrient levels<br>regulation of plasma lipoprotein levels | protein-lipid complex binding<br>response to lipoprotein particle<br>cellular response to lipoprotein stimulus<br>regulation of supramolecular fibre<br>cargo receptor activity | rhythmic process<br>gated channel activity<br>monoatomic ion channel activity<br>regulation of plasma lipoprotein levels<br>rhythmic behaviour |
| SS | **transcription coregulator binding**<br>rhythmic process<br>negative regulation of MAPK cascade<br>rhythmic behaviour<br>regulation of amide metabolic process | anatomical structure<br>homeostasis<br>production of inflammatory molecular mediator<br>**response to steroid hormone**<br>icosanoid metabolic process<br>**transcription coregulator binding** | monosaccharide metabolic process<br>sleep<br>**transcription coregulator binding**<br>rhythmic process<br>rhythmic behaviour |
| IS | **transcription factor coregulator binding**<br>rhythmic process<br>rhythmic behaviour<br>regulation of immune effector process<br>response to radiation | **transcription coregulator binding**<br>rhythmic process<br>rhythmic behaviour<br>**response to steroid hormone**<br>response to radiation | rhythmic behaviour<br>production of inflammatory molecular mediator<br>rhythmic process<br>icosanoid metabolic process<br>**transcription coregulator binding** |

## DISCUSSION

The objective of the analyses was not to criticize particular methods or approaches but rather create some clarity around strengths, weaknesses and pitfalls in this particular area of translational chronobiology. The findings show that the performance of single blood based multivariate biomarkers for melatonin phase, which is a proxy for SCN phase, is always much poorer when tested on independent validation sets compared to their performance on training sets. Performance depends on sample size of the training set, feature selection method and protocols from which training samples are derived. Performance of biomarkers developed on samples during which sleep occurs in phase with the melatonin rhythm is poor when applied to samples from protocols in which sleep is desynchronised from the melatonin rhythm. This desynchrony may occur in shift work, jet lag and some circadian rhythm sleep-wake disorders which are the typical use cases to which these biomarkers are supposed to be applied. Core clock genes do not feature prominently in the features selected to reflect SCN phase which implies that in the periphery they do indeed not reflect central circadian phase. Whether in the periphery they reflect local circadian rhythmicity or patency of circadian organisation remains an open question. Features selected by the transcriptomic-wide approaches vary greatly across methods and training protocols with only two elements of the glucocorticoid signalling pathway (*PER1* and *DDIT4)* being present in all predictor sets. *FKBP5* appeared in four of the predictor sets and is a chaperone protein that regulates activity of the glucocorticoid receptor. We have previously shown blood-based features that predict SCN phase were enriched for components of glucocorticoid signalling pathways (Laing et al., 2017). *PER3* also appeared in four predictor sets and has been shown consistently to have robust rhythmic expression in peripheral tissues (for review see Archer et al., 2018). Overall, these data show that the current omics and single sample-based biomarkers for SCN phase do not perform sufficiently well to meet the quantitative requirements of most research questions. Further application of well-established general principles for biomarker development and validation to the development of biomarkers for SCN phase and circadian rhythmicity can benefit the translation and application of basic circadian rhythm knowledge.

### Sample Size

Performance on the validation sets was poorer than the performance on the training sets for all sample sizes of the training set. The fact that performance of the biomarkers, when tested on a data set which is independent of the training set, improved with increasing sample size of the training set is not surprising. Development of models on small training sets are prone to overfitting. Overfitting as a cause of poor performance is very much related to the ratio of available features to sample size (Ng et al., 2023). Thus, also here, the main reason appears to be that the models over fit when the sample size of the training set is small. This explanation is supported by the observations that 1) the performance on the training set decreased when the sample size increased; 2) The discrepancy between performance on the training and validation sets decreased with increasing sample size. In some publications in the area of biomarker discovery, performance is not evaluated on an independent data set but assessed through cross validation. This evaluation method is, however, prone to high variance, as each model is tested on only a single sample, making performance estimates unstable and potentially misleading when assessing how the model will perform on new data (Austin et al., 2025).

### Comparing feature selection methods

Applying all methods to the same data sets and using the same performance metric (mean absolute error) when using one blood sample (rather than for example two samples, or a quasi-one sample, in which the sample is normalised relative to other samples of the same participant) allowed a direct comparison of the methods. The most obvious conclusion is that all methods performed rather poorly with mean absolute errors in the range of 3-4 hours when samples from a variety of protocols were included. There are to our knowledge no published guidelines for performance of biomarkers for SCN phase in humans. The required accuracy will very much depend on the application or use case. In the analyses presented here, which were all based on using only one sample, the absolute mean error was several hours, which is much larger than the estimated error of melatonin (Klerman et al., 2002). Although an error of several hours may be sufficient for some applications (i.e. delivering light exposure in the phase advance or phase delay region), it may be insufficient for other applications, i.e. determining whether onset of melatonin occurs before or after habitual bed time.

Overall PLSR and EN performed better than ZZ, but this better performance of these two methods was still rather poor. We note that we did not optimise the number of knots for the periodic smoothing splines of ZZ, but simply used the default. The performance based on a set of ‘clock genes’ was equally poor. The performance reported here is worse than the performance for these methods reported in a number of publications. This may in part be due to the performance metric used, the origin of the features (e.g., monocytes) or platform used (e.g., nanostring vs microarray or RNAseq) as well as the number of samples used in the training sets. Two other potential causes of this discrepancy are the validation approach, e.g., cross-validation vs independent data set, and the experimental conditions of the training set and validation set (see below). Please note that in our approach the stated purpose of the biomarker is the prediction of melatonin phase, whereas in some of the publications the purpose of the biomarker may have been detection of circadian phase in a particular tissue. How blood-based biomarkers may relate to tissue specific rhythmicity has been discussed elsewhere (Moller-Levet et al., 2022).

### Origin of training sets and validation sets: the impact of the presence, absence and timing of sleep

The results send a rather simple message. The more similar the training and validation sets with respect to the presence or absence of sleep (constant routine or a standard sleep-wake cycle) or timing of sleep (nocturnal sleep or daytime sleep), the better the performance of the biomarker. This is not surprising, but the important implication is that a biomarker developed from baseline data cannot be expected to perform well in, for example, a shift work situation or in sleep disorders such as delayed or advanced sleep wake phase disorder. Since biomarkers are used in those situations in which we don’t know circadian phase, it is best to draw training samples from a wide range of sleep-wake circadian paradigms or indeed from a target population. This recommendation is obviously not limited to blood-based biomarkers but extends to digital biomarkers for circadian phase based on large data sets from wearables and sometimes combined with mathematical modelling [see (Dijk and Duffy, 2020). These considerations also extend to blood-based biomarkers for sleep debt status [e.g., acute vs chronic sleep loss (Laing et al., 2019), and the development of algorithms to predict sleep stage in humans (Olesen et al., 2021)].

### Selected Features and Associated Processes

Biomarkers for circadian phase may, besides predicting circadian phase, also contain information on the nature of the target process. Here this relates to the features selected to predict melatonin phase, which is a proxy for SCN phase. Before we discuss the large variation in the features selected by the various methods and across the protocols, it is important to point out that the protocols used to develop and test the biomarkers did not have a major impact on the timing or amplitude of melatonin rhythm itself (see Archer et al., 2014 Fig 1; Archer et al., 2022 Fig 1; Lo et al., 2012 Supplemental Fig8). Furthermore, in all results presented here data were analysed with respect to the phase of the melatonin rhythm at which samples were collected. Thus, the melatonin rhythms under baseline conditions, constant routine conditions, mistimed sleep and constant bedrest conditions all had a similar amplitude. Likewise, for those protocols in which cortisol rhythmicity was assessed, no major changes were observed across conditions (see Archer et al., 2022; Bonmati-Carrion et al., 2024).

The feature sets constituting the biomarkers created by the various methods and across training protocols varied greatly with little overlap. The small overlap and the small number of ‘clock genes’ represented in these sets, may be surprising and should be explored. Given that the total number of features is approximately 40K, under baseline conditions 5-10% of the 40K features (i.e., 2-4K) are expected to be rhythmic, and the average predictor set contains far fewer features, it may be argued that the lack of overlap basically illustrates that many roads lead to Rome.

### The contribution of features and associated biological processes and molecular functions to predictor sets

Core clock genes were not enriched as features in the training sets used here and when a specific set of clock genes was used for training with PLSR it generally performed worse than predictors from the unbiased training sets. However, some clock genes do consistently feature as predictors across multiple studies. *PER1*, which featured in all training sets for all methods in this study, has also been identified in four previous studies (Hughey, 2017; Laing et al., 2017; Braun et al., 2018; Wittenbrink et al., 2018). *PER1* expression is induced by glucocorticoid (Yamamoto et al., 2005 and is less regulated by the sleep/wake cycle (Mongrain et al., 2010) unlike *PER2* which in the periphery is significantly affected by the sleep-wake cycle (Mongrain et al., 2010; Hoekstra et al., 2021). This could underlie PER1’s prominence as a detection feature in human blood taken from a variety of sleep/wake conditions. *DDIT4* (also known as *REDD1*) also featured in all training predictor sets in all methods. DDIT4 is involved in stress response signalling, is regulated by glucocorticoid (Zhidkova et al., 2022; Choi et al., 2024) and regulates glucocorticoid receptor function (Chudakova et al., 2023). Expression of *FKBP5* is strongly induced by glucocorticoid (Martinelli et al., 2024) and its coded protein is a chaperone that interacts with the glucocorticoid receptor and regulates its activity (Fries et al., 2017). *FKBP5* was a predictor in four training sets in all three methods and it (or its paralog *FKBP4*) has featured in four previous studies (Hughey, 2017; Laing et al., 2017; Wittenbrink et al., 2018; Wu et al., 2018). Recent data have shown that overexpression of *Fkbp5* in mouse forebrain enhanced circadian amplitude and decreased rhythm fragmentation (Gebru et al., 2024). Together, these results point towards the significant contribution of elements of peripheral glucocorticoid signalling pathways in models developed to predict central circadian phase. For a summary of recent studies that have developed transcriptomic predictors for circadian phase see Table S4.

Although core clock genes did not feature significantly as phase predictors in this study, the GO terms ‘rhythmic process’ and ‘rhythmic behaviour’ were enriched among the predictor sets in each method. Other enriched GO terms that occurred across multiple methods included ‘response to steroid hormone’ and ‘transcription coregulator binding’. These GO terms also emphasise the contribution of glucocorticoid signalling elements. Apart from these, other GO terms for biological processes and molecular functions differed across both training set and method highlighting the fact that different training sets produce predictor sets with different features. However, it should also be noted that a variety of GO terms associated with lipid metabolism did occur across training sets and methods. While identified biomarker transcripts predictive for circadian phase cannot be directly correlated with specific biological functions and molecular processes, their enrichment within such gene ontology terms does provide some indication of processes and functions that robustly contribute to peripheral circadian phase.

### General Conclusion

Given that the human circadian timing system is a hierarchical multi oscillator system in which rhythmicity is generated by a multitude of feedback loops at the level of transcription and translation modulated by systemic and behavioural factors, identification of specific biomarkers will be challenging. This applies to biomarkers based on physiological, endocrine, transcriptomic, metabolomic and proteomic features. Here we have focussed on the simple case of transcriptome-based biomarkers for SCN phase. The analyses demonstrate that the impact of behaviour, i.e. the sleep-wake cycle and associated factors, on the transcriptome makes it challenging to design biomarkers which reliably indicate SCN phase under conditions most relevant to use cases such as shiftwork and jetlag. These considerations/concerns apply independent of the methos used for feature selection.

## Acknowledgments

DJD is supported by the UK Dementia Research Institute [award number CF2023\7-UKDRI-7206.] through UK DRI Ltd, principally funded by the UK Medical Research Council, and additional funding partner Alzheimer’s Society, and the National Institute for Health Research (NIHR) Oxford Health Biomedical Research Centre (BRC), (NIHR203316). The study was funded by grants from the Biotechnology & Biological Sciences Research Council BB/N004981/1 BB/F022883, and the Air Force Office of Scientific Research FA9550-08-1-0080. The study was supported by the European Space Agency (ESA), the UK Space Agency (UKSA), and the National Institute for Health and Care Research (NIHR) Oxford Health Biomedical Research Centre. The views expressed are those of the author(s) and not necessarily those of the NIHR or the Department of Health and Social Care (NIHR Oxford Health Biomedical Research Centre grant reference number: NIHR203316). The authors would like to acknowledge contributions from Emma E Laing, Colin Smith, Giselda Bucca, Malcolm von Schantz, Maria Bonmati-Carrion, Nayantara Santhi, Giuseppe Atzori, Sylvia Kaduk, Jeewaka Mendis and members of the Surrey Clinical Research Facility. We thank Benita Midleton for analysing the melatonin samples, and the staff at the MEDES clinic in Toulouse for the bed rest study.

## Author Contributions

CM-L, SNA and DJD conceived the study, CM-L, SNA and DJD analysed the data, CM-L, SNA and DJD prepared figures/tables, all authors wrote and reviewed the manuscript.

## Statements and Declarations

The authors declare no conflicts of interest related to the study.

## Ethical Considerations

All studies reported here were conducted in accordance with the Declaration of Helsinki. The sleep restriction study and the mistimed sleep study received a favourable ethical opinion from the University of Surrey Ethics Committee. The bed rest study was approved by the Comite de Protection des Personnes SUD-OUEST ET OUTRE_MER 1.

## Consent to Participate

All participants involved in the studies reported here provided informed written consent.

