## Supplemental Material for "Performance of blood-based biomarkers for human circadian pacemaker phase: Training sets matter as much as feature selection methods"

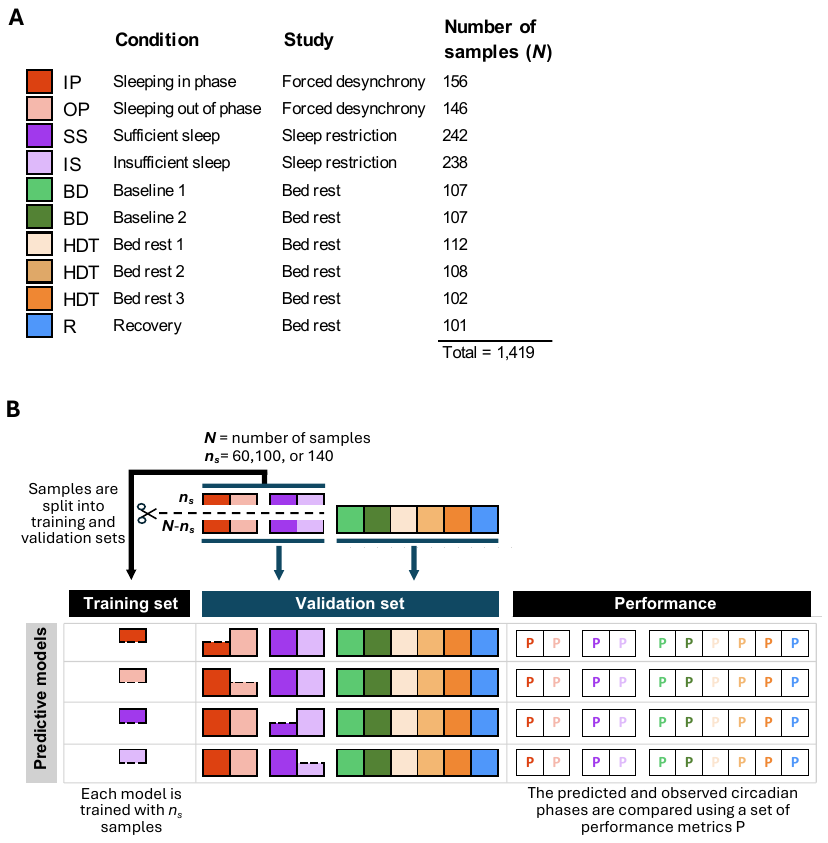

**Supplemental Figure 1. Methodological approach. A** Description of experimental conditions and number of samples. **B** Diagram of methodological approach. Training sets were created by randomly selecting *n_s_* samples from the total *N* samples of condition *X*. We used two selection strategies: one in which samples were randomly selected, and another in which samples were also randomly selected, but with no overlap between participants in the training and validation sets. The validation set included the remaining (*N-n_s_*) samples from condition *X*, along with all samples from other conditions. A predictive model was developed on each training set and used to predict circadian phase in the validation set. The predicted and observed circadian phase were compared, and a set of performance metrics (P) were calculated independently for each experimental condition in the validation set.

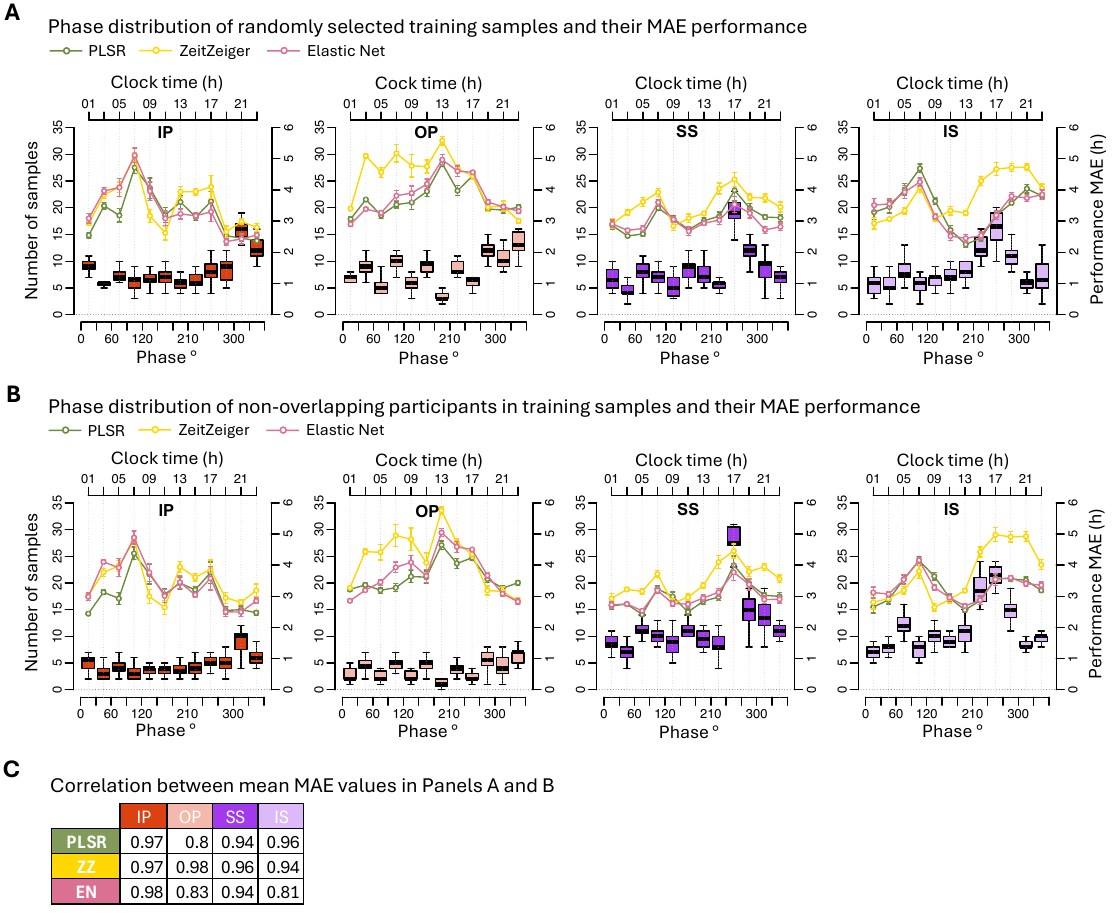

**Supplemental Figure 2. Effect of training sample selection strategy on phase distribution and prediction performance across the circadian cycle.** **Panel A** shows results from training sets composed of randomly selected samples (with possible participant overlap), while **panel B** shows results from training sets constructed using non-overlapping participants. For each of the four training conditions, the number of training samples (boxplots, left y-axis) and prediction performance (mean absolute error [MAE], line plots, right y-axis) are shown as a function of circadian phase (°, bottom axis) and clock time (h, top axis). Boxplots represent the distribution of sample counts across 30° phase bins from 20 random draws of 100 samples. MAE values for three prediction methods - PLSR (green), ZeitZeiger (yellow), and Elastic Net (pink) - reflect the average MAE across the four validation sets, with error bars indicating standard deviation. Panel **C** presents the Pearson correlation between the MAE values in panels A and B (n=12 per cell). All Benjamini–Hochberg (BH)-corrected correlation p-values were < 0.005. In all panels, columns correspond to the conditions used to derive the training sets. IP = sleeping in phase during the night, OP = sleeping out of phase during the day, SS = constant wake after one week of sufficient sleep, IS = constant wake after one week of insufficient sleep.

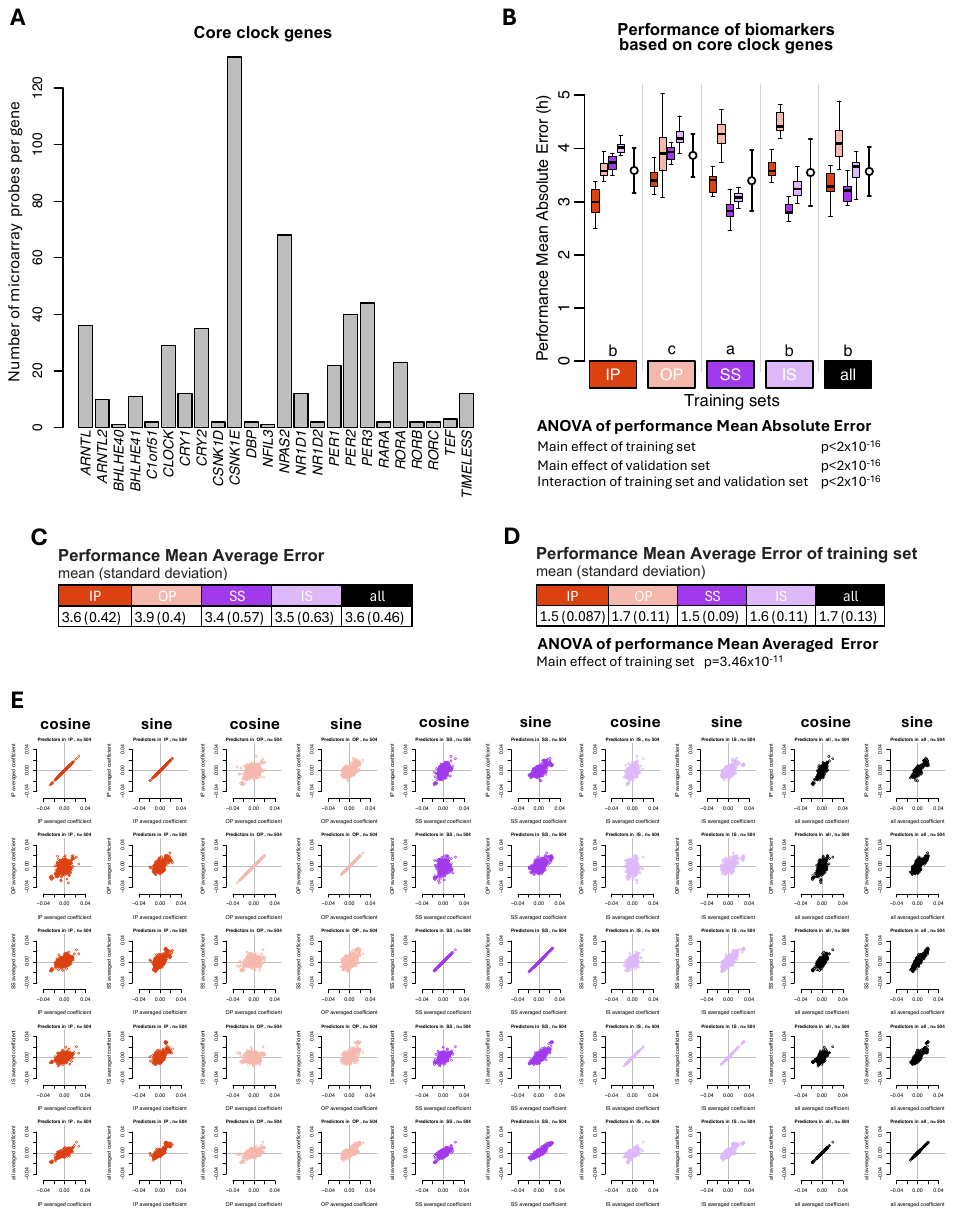

**Supplemental Figure 3. PLSR based on a fix set of clock genes**. **A** 24 core clock genes were mapped to 504 microarray probes and used as predictors in the generation of PLSR biomarkers. **B** Performance results obtained for PLSR. Twenty training sets of 100 samples were generated for each experimental condition. Each coloured box plot represents the distribution of 20 mean absolute error (MAE) values for each condition in the validation set, with outliers omitted for clarity. White open circles with error bars indicate the mean and standard deviation of all 80 validation values per training set (20 sets x 4 conditions). The two-way ANOVA for MAE showed significant effects of both the training set and validation set as well as their interaction (all p < 2×10⁻¹⁶). The Compact Letter Display (CLD) summarizes the results of pairwise comparisons between different training sets, as determined by Tukey’s test conducted on the two-way ANOVA of training and validation sets. **C** Mean and standard deviation of 80 MAE values per training condition (20 training sets x 4 validation conditions) shown in panel **A**. **D** Mean and standard deviation of 20 MAE values per training condition (20 training sets per condition), where the same samples are used for both training and validation. The ANOVA for MAE of the training set showed significant effect of training set (p=3.46×10^-11^). In all panels, training sets are depicted as coloured rectangles on the horizontal axis. **E** Averaged regression coefficients (across the 20 runs) of the 504 predictor transcripts in the models generated in panel **A**. IP = sleeping in phase during the night, OP = sleeping out of phase during the day, SS = constant wake after one week of sufficient sleep, IS = constant wake after one week of insufficient sleep. The condition “all” includes an equal number of samples from each of the four experimental conditions. IP = sleeping in phase during the night, OP = sleeping out of phase during the day, SS = constant wake after one week of sufficient sleep, IS = constant wake after one week of insufficient sleep.

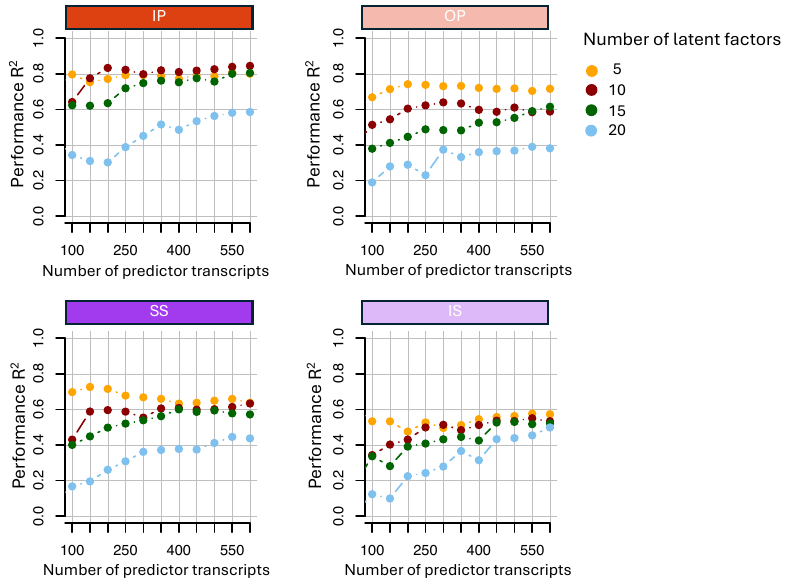

**Supplemental Figure 4. Selection of number of predictor transcripts and latent factors for PLSR models.** Leave-one-participant-out cross-validation performance R^2^ applied to training set when using different combinations of number of predictor transcripts (horizontal axis) and latent factors (coloured circles). Each panal corresponds to a different training set condition. All training sets were restricted to 140 samples.

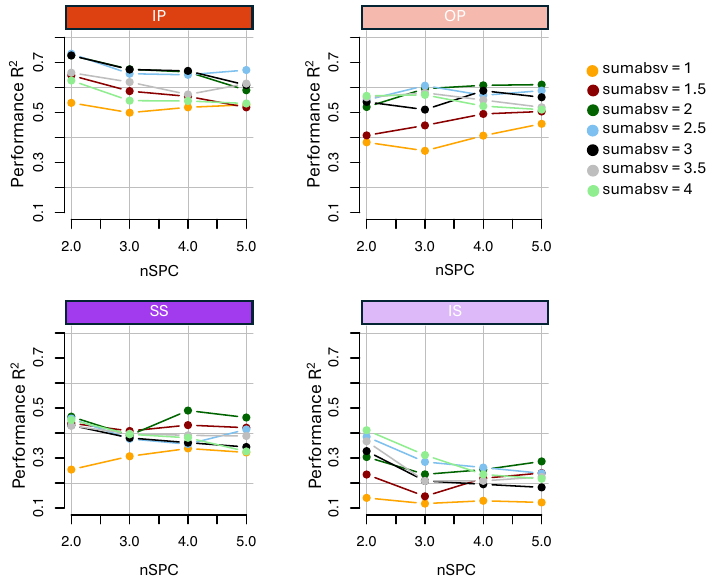

**Supplemental Figure 5. Selection of nSPC and sumabsv values for ZZ models.** Leave-one-participant-out cross-validation performance R2 applied to training set when using different combinations of nSPC (horizontal axis) and sumabsv (coloured circles) values. Each penal corresponds to a different training set condition. All training sets were restricted to 140 samples.

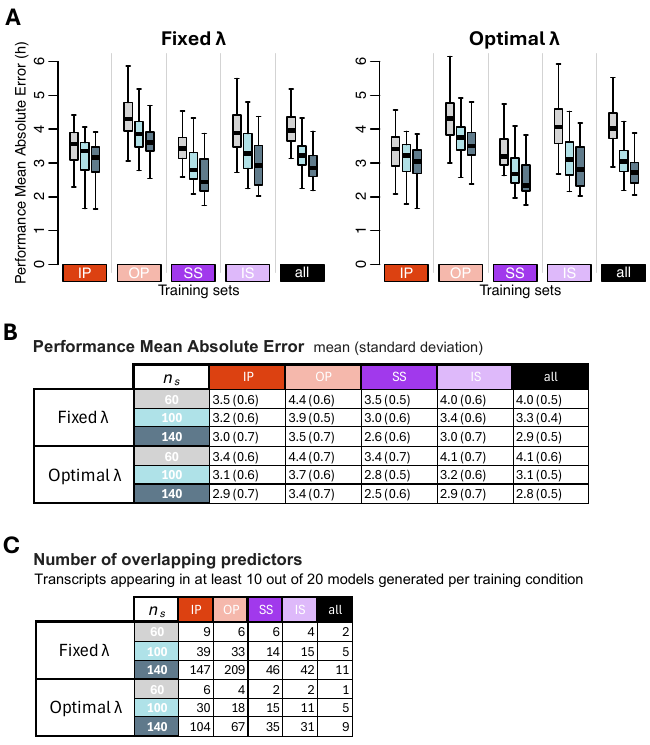

**Supplemental Figure 6. Effect of lambda selection (fixed vs. optimised) on EN model performance.** **A** Performance results obtained for two lambda selection strategies: Fixed (left) and Optimal (right). Each box plot represents the distribution of 20 mean absolute error (MAE) values for each condition of the validation set, totalling 80 values (20 sets x 4 conditions), with outliers omitted for clarity. **B** Mean and standard deviation of 80 MAE values per training condition (20 training sets x 4 validation conditions) shown in panel **A**. **C** Number of overlapping predictors, defined as transcripts appearing in at least 10 out of the 20 models generated per training set. In each panel, results were obtained for *n_s_* values of 60 (gray box plots), 100 (light blue box plots) and 140 (dark blue box plots), and training sets are represented by coloured rectangles on the horizontal axis. The training condition “all” includes an equal number of samples from each of the four experimental conditions. IP = sleeping in phase during the night, OP = sleeping out of phase during the day, SS = constant wake after one week of sufficient sleep, IS = constant wake after one week of insufficient sleep.

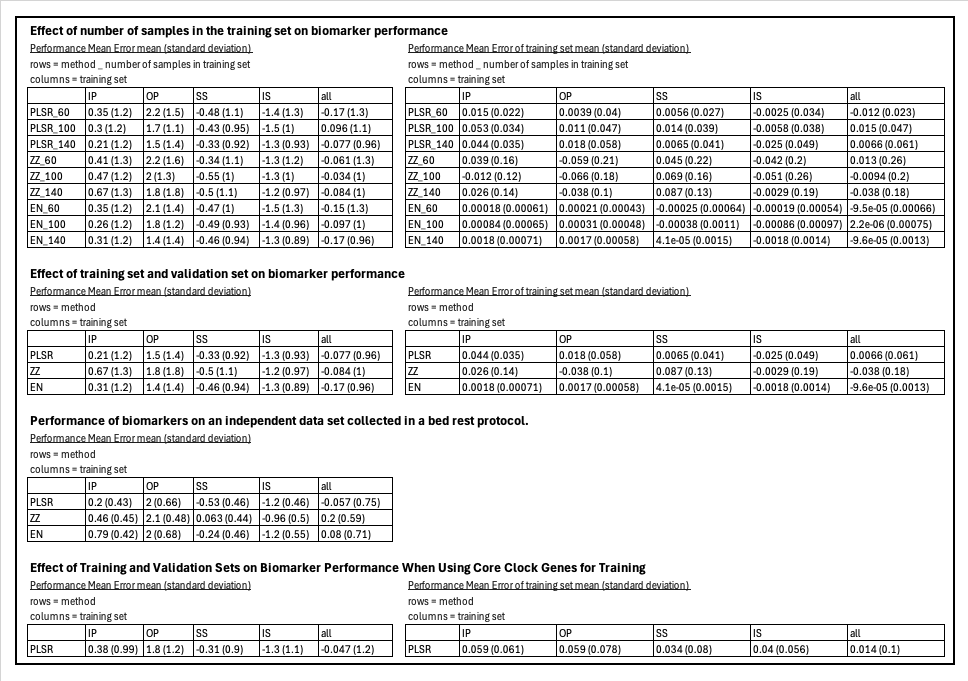

**Table S1.** Performance results based on Mean Error

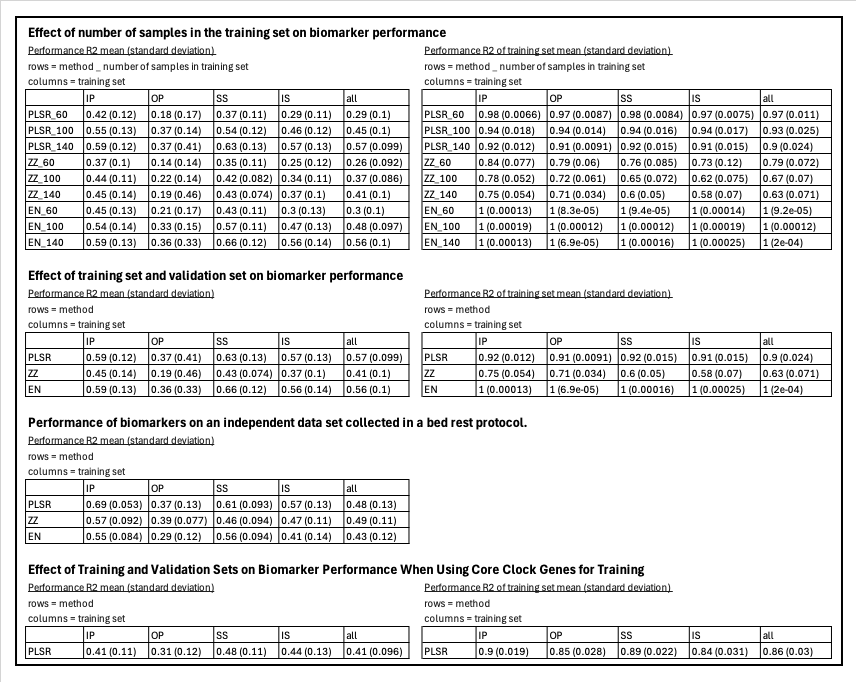

**Table S2.** Performance results based on R^2^

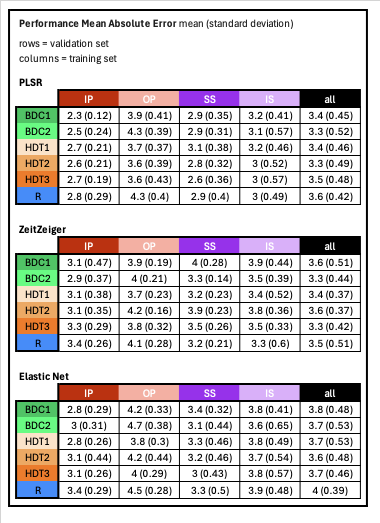

**Table S3.** Performance based on an independent dataset

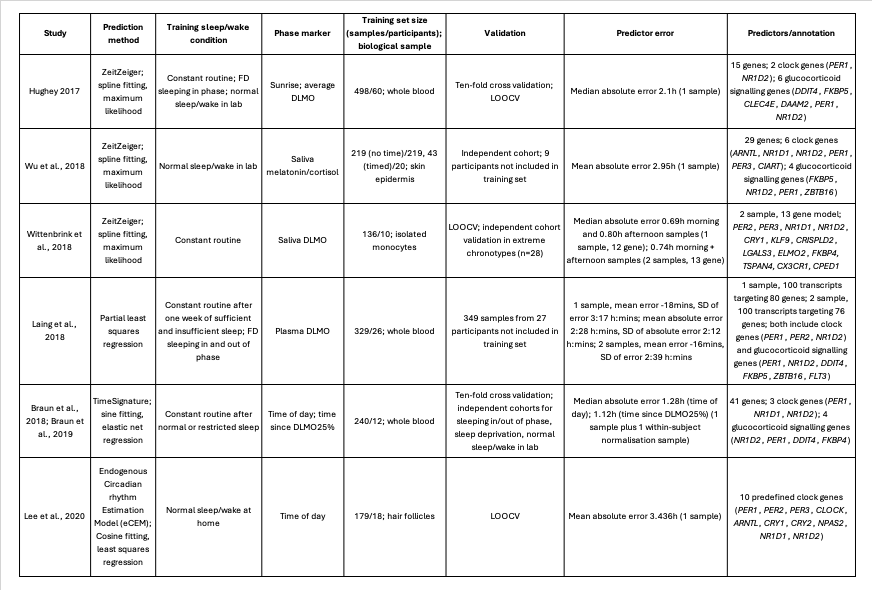

**Table S4.** Comparison of studies that have used different approaches to develop transcriptomic predictors of circadian timing.
